# Biophysical modeling of thalamic reticular nucleus subpopulations and their differential contribution to network dynamics

**DOI:** 10.1101/2024.12.08.627399

**Authors:** Polina Litvak, Nolan D. Hartley, Ryan Kast, Guoping Feng, Zhanyan Fu, Alexis Arnaudon, Sean L. Hill

**Affiliations:** Blue Brain Project, École polytechnique fédérale de Lausanne (EPFL), Campus Biotech, Geneva, Switzerland; Stanley Center for Psychiatric Research, Broad Institute of MIT and Harvard, Cambridge, MA 02142, USA

## Abstract

The burst firing mode of thalamic reticular neurons plays a pivotal role in the generation and maintenance of sleep rhythms and is implicated in sleep-related deficits characteristic of neurodevelopmental disorders. Although several models of reticular neurons have been developed to date, we currently lack a biophysically detailed model able to accurately reproduce the heterogeneity of burst firing observed experimentally. Using electrophysiology recordings of patch-clamped fluorescently tagged Spp1+ and Ecel1+ reticular neurons, we leverage a previously established statistical framework to introduce differentiation of cell types in model thalamic reticular neurons. We developed a population of biophysically detailed models of thalamic reticular neurons that capture the diversity of their firing properties, particularly their ability to generate rebound bursts. These models incorporate key ion channels, such as T-type Ca^2+^ and small conductance potassium channels (SK), and enable systematic investigations into the impact of these channels on single-cell dynamics. By integrating these models into a thalamic microcircuit, we demonstrate that T-type Ca^2+^ and SK channel conductances have opposing effects on spindle oscillations. We identify a simple relationship between these conductances and the peak firing frequency of spindles, maintained across circuits with mixed reticular neuron populations, providing a framework for understanding how ion channel expression influences thalamic network dynamics. Collectively, these models establish a foundation for relating intrinsic cellular properties of reticular cell populations to network-level activity in both healthy and pathological conditions.

## Introduction

The thalamic reticular nucleus (TRN) is a shell-like subcortical structure surrounding the dorsal thalamus. It is composed of a heterogeneous population of gamma-aminobutyric acid (GABA)-ergic neurons and provides a major source of inhibition to the thalamus in rodents [Pinault, 2004]. Upon stimulation, reticular neurons generate action potentials (APs) in two distinct firing modes, tonic and burst, depending on their membrane potential and their expression of low-threshold voltage-gated T-type Ca^2+^ channels [Contreras D, 1993]. At resting or relatively depolarized membrane potentials, reticular neurons fire regular tonic sodium spikes. At hyperpolarized potentials, they generate repetitive low-threshold calcium transients superimposed with high-frequency sodium spikes, referred to as bursts. TRN bursts are efficient at producing postsynaptic inhibition, while rebound bursting in both TRN and thalamic relay cells plays a crucial role in generating and maintaining thalamocortical rhythm [Steriade M, 1993]. Multiple previous studies establish the reticular thalamus as the origin of spindle activity, a major brain oscillation most prevalent in non-rapid eye movement (NREM) sleep, further regulated by thalamocortical interactions [Destexhe Alain, 2007, McCormick and Bal, 1997, Steriade M, 1993]. In naturally sleeping rodents, the duration and frequency of spindle oscillations are shown to be directly shaped by reticular inhibition [Bartho et al., 2014]. Recently, a computational thalamoreticular circuit model was able to link spindle duration to membrane potential dynamics in both thalamic relay and reticular neurons [Iavarone et al., 2023]. The distinct firing properties of TRN neurons critically depend on the subtypes of ion channels they express. In particular, a large body of literature identifies the T-type low threshold calcium channel family as a mediator of burst firing in these neurons [Contreras D, 1993, Li et al., 2020]. What is less well understood is which subtypes expressed in TRN are necessary for rhythm generation and what their respective contributions are to the differential propensity of reticular cells to rebound burst. Furthermore, TRN bursts are typically followed by an afterhyperpolarization (AHP) generated by small-conductance calcium-activated (SK)-type potassium currents. How T-type Ca^2+^ channels and SK channels work in concert to shape rebound burst activity in TRN cell subpopulations is one of the aspects addressed in this work. TRN cells greatly vary in their propensity to generate bursts [Clemente-Perez et al., 2017, Contreras D, 1993, Lee SH, 2007]. Two recent studies, using immunohistochemical staining for molecular markers [Martinez-Garcia et al., 2020] as well as transcriptomics [Li et al., 2020] give an unprecedented characterization of reticular cells in terms of cellular morphology, molecular expression, axonal connectivity, and physiological activity. According to their multiscale single-cell analysis, the heterogeneity of TRN cells is characterized by a transcriptomic gradient composed of two negatively correlated gene expression profiles. Neurons at the extremes of the gradient have a near-exclusive expression of a few marker genes used to segregate them into a core (Spp1+) and shell (Ecel1+) TRN subpopulation with a differential propensity to generate intrinsic rebound bursting [Li et al., 2020] and consequently a differential contribution to network dynamics.

Although several computational models of reticular neurons have been developed to date [Destexhe et al., 1994, McCormick and Huguenard, 1992], a biophysically accurate model capable of effectively capturing the diverse continuum of TRN electrical properties and replicating the heterogeneity of TRN rebound burst firing remained unavailable. Using data obtained from novel Cre mouse lines targeting genetically segregated TRN populations expressing the Spp1 and Ecel1 genes [Hartley et al., 2024], and leveraging the Markov chain Monte Carlo (MCMC) approach of Arnaudon et al. [2023], we built electrical reticular models that reproduce the full range of firing modes observed experimentally. We then incorporated these models into the thalamoreticlar microcircuit of Iavarone et al. [2023], delineating the differential contributions of TRN intrinsic conductances to spindle-like activity as well as predicting the relative fraction of Ecel1+ and Spp1+ in the somatosensory TRN from spindle properties under knock-down experiments of Ca^2+^ channels similar to [Li et al., 2020]. Overall, this modeling work provides important insights into how the intrinsic cellular properties of reticular cells shape emergent circuit dynamics, advancing our understanding of thalamic rhythmogenesis in both health and disease.

## Results

### Experimental data

This modeling work is based on electrophysiology data of Hartley et al. [2024], recorded from two transcriptomically defined populations of TRN neurons, referred to as Ecel1+ and Spp1+, which represent the two genes most deferentially expressed by the identified genetic profiles. The dataset consists of whole-cell patch clamp recordings from 52 genetically labeled Spp1+ or Ecel1+ neurons, subjected to several depolarizing and hyperpolarizing current step protocols. For the protocol assessing rebound bursting activity, referred to as the burst protocol, neurons were subjected to hyperpolarizing current injections to a voltage near maximal de-inactivation of T-type Ca^2+^ channels to induce maximal rebound bursting for each cell, starting from various holding potentials. On average, Spp1+ neurons displayed a larger number of bursts than Ecel1+ neurons, and for a subset of holding voltages, roughly half of the Spp1+ neurons were able to continue bursting past the 15 second sampling window (and up to several minutes) and were therefore defined as Spp1+ runaway bursting neurons Hartley et al. [2024]. Consequently, each cell from the experimental database was classified into one of three electrical types Ecel1+, Spp1+ or Spp1+ runaway (referred to as runaway). We display some voltage traces of these recordings in Fig. 1a with a zoom on the first burst in Fig. 1b, showing some variability in the number of APs per burst. The entire scan of initial holding voltages of the burst protocol is shown Fig. 1c for one cell per type, which we will call the burst curve of the cell. We fit a skewed Gaussian function (see Method and Fig. 8) to these points to extract important parameters such as curve centre location in millivolts (mV), amplitude, and skewness of the burst curve. In Fig. 1d-e we show the distribution of two of these parameters that have statistical differences across cell types (see Fig. 8 for the others). As expected, the burst curve amplitude is largest for runaway and smallest for Ecel1+ cells, but interestingly, burst curves for Ecel1+ are peaking at higher voltages than Spp1+. This is consistent with the fact that they express more of the Cav3.1 T-type Ca^2+^ channel subtype (see Fig. 15), known to open at higher voltages [McRory et al., 2001] than the other subtypes. These burst curves have large variability even within the same cell type. In Supplementary information (SI) Fig. 8, we present a broader range of burst curves, highlighting the variability in their shapes, including cells with highly skewed curves or curves lacking a distinct peak.

**Figure 1:**
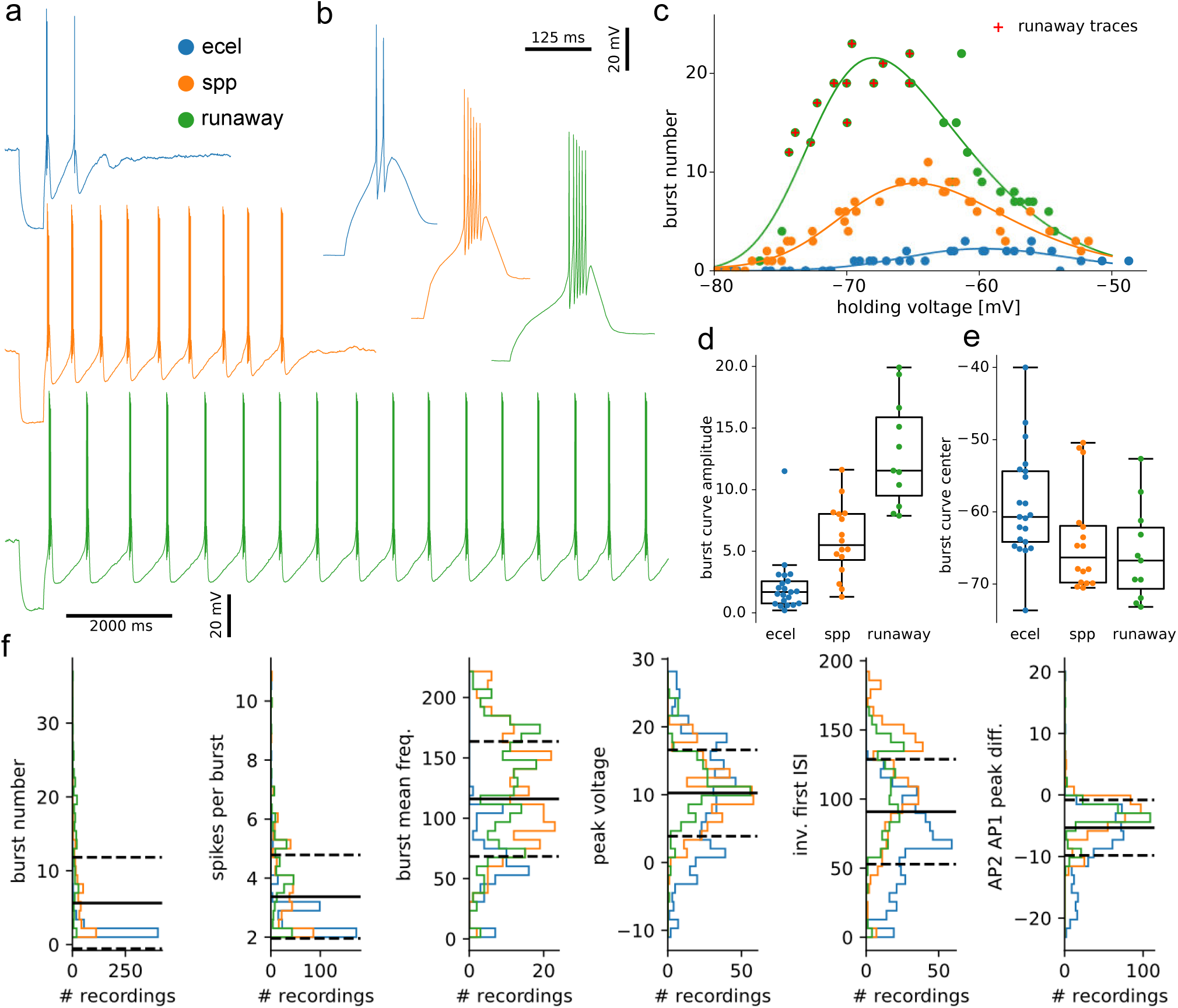
Experimental data of thalamic bursting cells. **a.** Exemplar traces for Ecel1+ (top), Spp1+ (middle) and runaway (bottom). **b.** Zoom on the first burst of each trace from panel a. **c.** Burst number as a function of holding voltage (burst curves) for Ecel1+ (left), Spp1+ (middle), runaway (right) of the cells of panel a. Red cross indicates traces with runaway property, the line is a fit of a skewed Gaussian curve (see Method). **d.** Distribution of amplitude parameter of the skewed Gaussian fit of each cell per cell type. **e.** Distribution of centre parameter of the skewed Gaussian fit of each cell per cell type. **f.** Distribution of the main electrical features used for model building, split by cell type. The black line is the average of combined data across three cell classes and the dashed line represents the first standard deviation.

In addition to bursting properties, we summarize the important features of the experimental data used to build our models in Fig. 1f, for each cell type. This data contains all recordings with bursts, not just traces with maximum elicited bursts per cell (like the burst curves), and shows that many features are similar across cell types. It is worth noting that the first inter-spike interval (ISI) is shorter for Spp1+/Runaway cells than for Ecel1+ and is correlated with more action potentials per burst, as seen from Fig. 1b and SI Fig. 13. As we will see later, there is a significant correlation between the intra-burst AP frequencies, first ISI and the number of APs per burst, which can be related to the amount of SK current.

To further quantify the extent to which the extracted electrical features represent these putative cell types, we employed an XGBoost feature classifier [Chen and Guestrin, 2016] to classify cells based solely on their feature values (see SI Fig. 10). We obtained a classification accuracy of 89 ± 14.7% when using all extracted electrical features, and only a marginal drop of around 1% when omitting the burst number, showing that electrical features such as the runaway measure (tracking slow AHP depth between bursts), burst frequency or time to first spikes (top three most important features) are already well representative of these differences in intrinsic electrophysiological properties between the three cell types.

### Building electrical models of bursting cells with Markov chain Monte Carlo

To build and study the full range of electrical models of TRN cells, and in particular, the three subtypes identified experimentally, we use the Markov chain Monte Carlo (MCMC) approach from our prior work Arnaudon et al. [2023]. Instead of a direct optimization such as was done previously [ Iavarone et al. [2019], Van Geit et al. [2016]], we sample many electrical models from a probability distribution inversely proportional to a cost function, constructed by comparing electrical features of the model with experimental data. This allows us to create many models with a low value of the cost function encompassing all subtypes we are interested in, making it possible to understand the differences between them. In particular, the distinction between Ecel1+ and Spp1+ cells is a continuum rather than a dichotomy Li et al. [2020], hence creating many models spanning the full range of electrical features obtained experimentally, rather than building one specific model per cell type, provides a more detailed understanding of these TRN cell subpopulations. In more detail, the cost function is constructed from the z-scores of a set of electrical features extracted from traces under specific protocols of somatic current injections. The z-scores are computed against mean and standard deviations, adjusted from experimental recordings to encompass all electrical subtypes. Some features, such as the burst number (see Method for its definition), have been set to a large standard deviation to ensure they do not constrain the model too strongly (see Table 1 for all parameters). In principle, these values should be extracted from experimental data [Arnaudon et al., 2023, Reva et al., 2023, Van Geit et al., 2016], but in the present case, the recordings were obtained from various holding voltages, and due to the large variability of the burst curves, it is impossible to design a consistent automated feature extraction method. In addition, using the entire burst curve would require the numerical evaluation of many traces and be too computationally expensive. We therefore used features from three versions of the same protocol. The protocol employed is comprised of the following three current steps: 1) a holding step to maintain the model at a target voltage, 2) a hyperpolarizing step to bring the voltage to −100 mV, 3) a depolarizing step to return the model to the initial holding voltage. We inject the corresponding current via a bisection search and a step protocol measuring the voltage after it stabilizes to target a specific holding voltage. The three variants of this protocol differ between the holding voltage: −80mV, −65mV and −55mV. The first target value ensures the cell does not burst after the hyperpolarization step, while the next two ensure it does. The third protocol is not strictly necessary but was included to force the cell to burst for holding voltages up to at least −55mV and to be able to compare the number of elicited bursts at two different voltages, to for example assess if the burst curve exhibits a peak.

The electrical models we implemented are a refined version of the model of Destexhe et al. [1994], similar to the thalamic relay cell models of Iavarone et al. [2019] or more recently the TRN models of Iavarone et al. [2023]. We used a detailed morphology (with only soma and basal dendrites) taken as the average morphology from the population of experimental reconstructions used in Iavarone et al. [2023] (see SI Fig. 9 and Fig. 2a for an illustration of the morphology). We placed Na and K channels in the soma to generate action potentials (hh2 Na and hh2 K), generic T-type Ca^2+^ channels (I_T_), and generic SK channels (I_AHP_) in both soma and basal dendrites to generate low-threshold Ca^2+^ spikes. Ca dynamics (cad) with ATPase pump and first-order buffering is also included in all compartments. We also included Ca^2+^-dependent nonspecific cation current (I_CAN_) and fast transient potassium current (I_A_) in basal dendrites to control rebound burst properties. In addition, in line with experimental reports of higher density of T-type Ca^2+^ channels in the distal dendrites of TRN [Crandall et al., 2010], we implemented a linear increase of I_T_ conductance in basal dendrites, parametrized by a slope in addition to the maximum conductance (see Method and Table 2 for more details on the model). We also considered a parameter to apply a voltage shift of the T-type channel mechanisms, to control the location of the current window of this ion channel, known to be important for bursting properties. We will show later how this shift can be related to relative conductances of Cav3 subtypes.

**Figure 2:**
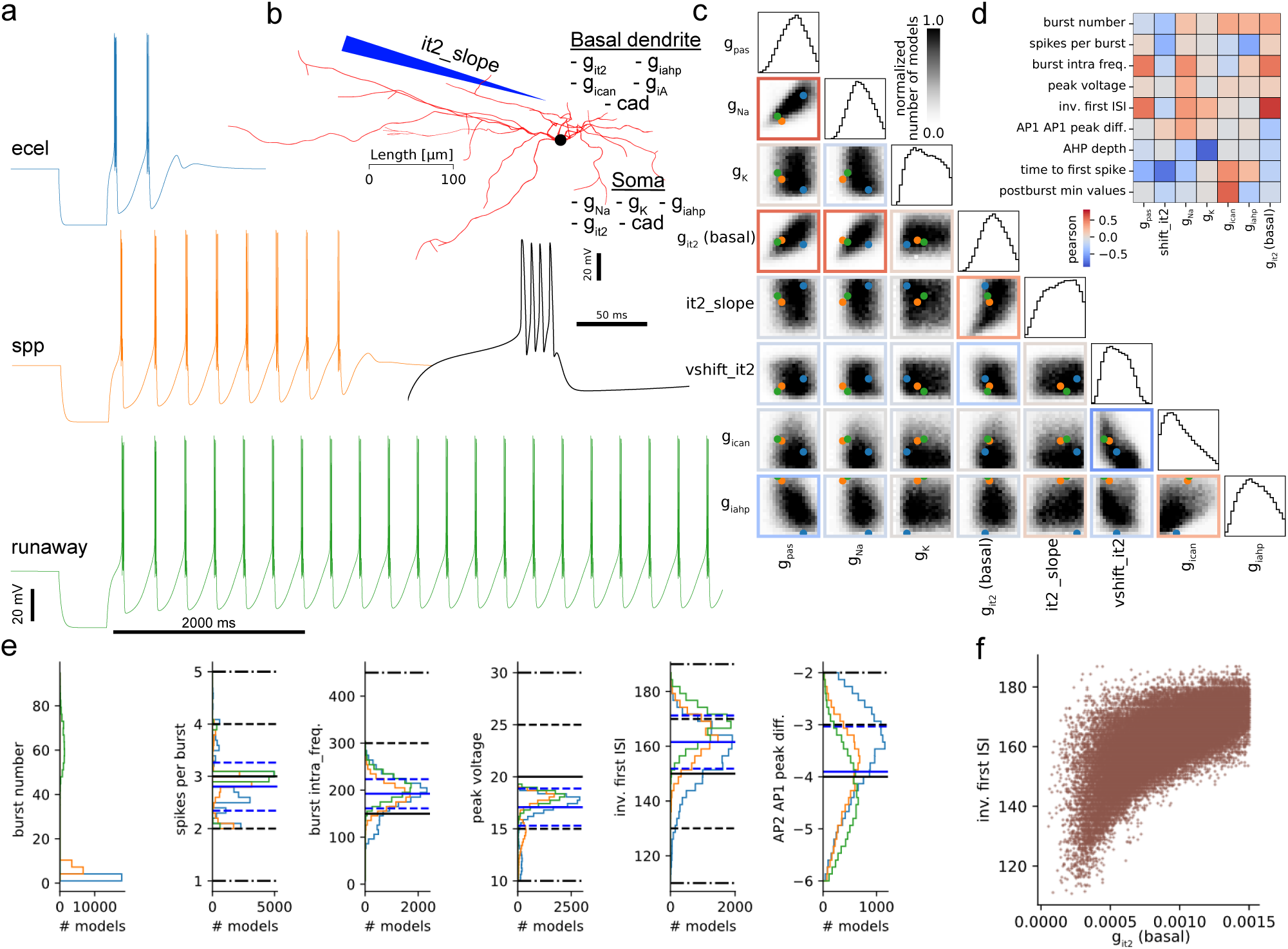
Model building of four types of TRN cells. **a.** Three traces of corresponding models of each electrical type. Zoom on the first burst of Ecel1+ model in the inset. **b.** Detailed reconstruction of the morphology used for model building, with a list of ion channels in each compartment (see Method and SI Fig. 9). **c.** Corner plot of densities of models with std*<* 2. Frames are colored with Pearson correlation between parameters, dots represent the location of the three models of panel a. **d.** Pearson correlations between some model parameters and features. **e.** Feature distribution of the main features used for MCMC constraints, split by cell type. The mean (black line), 1 std (dashed line) and 2 std (dot-dashed lines) are the constraints used in the cost function (see SI Table 1). Blue lines are the mean and 1 std of the data, with all cell types combined. **f** Scatter plot of inverse first ISI vs T-type Ca^2+^ (I_T_) conductance, a positively correlated pair of parameter/feature, as panel d.

We run MCMC with bounds of model parameters adjusted to capture a large part of the parameter space that contains valid models, defined as models with maximal scores less than 2 standard deviations. These bounds are manually adjusted from trial and error until a satisfactory solution is reached in terms of features and from visual inspections of traces. In Fig. 2c, the corner plot of model densities in the parameter space shows significant correlations between many pairs of parameters (colored frames with Pearson correlations). For example, I_T_ conductance in basal dendrites is positively correlated with g pas and somatic Na, due to the constraint imposed by experimental data on the relative amplitude of the first two action potentials (not shown). Indeed, for larger low-threshold Ca^2+^ spikes in the basal dendrites, the soma requires more Na to trigger Na spikes at a fast rate and a similar amplitude. The slope of the I_T_ distribution in basal dendrites and its maximal conductance are also positively correlated, as for larger it2 slope values (smaller slope), the total amount of I_T_ is reduced, hence compensated with a global increase via the I_T_ conductance parameter. On the contrary, I_CAN_ conductance and the voltage shift parameter of the T-type channel mechanism (see Method) as well as I_AHP_ and passive conductances have a negative correlation from the constraint imposed by the slow AHP depth and burst intra-frequency features (these features become large away from the sampled space of models at 2 std, not shown).

From the many models we obtained, we show three representatives of each cell class under the rebound burst protocol at −65*mV* in Fig. 2a, similar to the experimental data of Fig. 1a. The parameters for these randomly chosen models are shown with colored dots in Fig. 2c and in SI Table 2. For this particular set of models, the primary distinction between Ecel1 and Spp1/runaway cell types can be attributed to significant differences in I_AHP_ conductance (see also Fig. 14). In contrast, the parameters of Spp1 and runaway models exhibit greater similarity.

In Fig. 2d, we present Pearson correlations between model parameters and electrical features, thereby identifying the parameters that exert the most significant influence on each feature. For example, I_T_ conductance in basal dendrites controls the first ISI (and the burst mean frequency, as ISI are fairly regular) shown in more detail in the scatter plot in Fig. 2f. Indeed, a larger amount of I_T_ in basal dendrites will generate larger low-threshold spikes [Cain and Snutch, 2010], creating stronger bursts of Na spikes in the soma, with more and higher frequency action potentials.

Additionally, we analyze correlations between experimental and model features summarized in Fig. 13. Our models recover the most prominent correlations between the number of APs in a burst, intra-burst frequency and first inter-spike interval (ISI). Due to the high variability of experimental burst curves, and that the experimental features are based on mean values gathered from recordings obtained from many different holding voltages, several correlations do not match. Additionally, our models are built with generic ionic mechanisms and hence overlook the nuanced differences known to exist between ion channel isoforms deferentially expressed by Ecel1+ and Spp1+ neurons (see SI Fig 15).

In Fig. 2e, we show the distributions of the main features used by MCMC, the mean and std we used to build the cost function shown in black (see SI Table 1) and the mean and std of all the samples with std*<* 2 highlighted in blue. We can reproduce the difference in first ISI between cell types, as seen in the experimental data across a large range of holding voltages (see Fig. 1f), or the average number of spikes per burst. The action potential (peak voltage) amplitude feature in the experimental data has a large variability, we therefore chose to restrict it to high values with a mean value at 20*mV*, as modeling this variability is out of the scope of this work. For the relative amplitude of the first two APs, we see an opposite trend to experimental data, with Ecel1 models having a smaller decrease than the other two model types. Interestingly, the drop is larger in the experimental data, even excluding the long tail between −10 and −20mV, with cells able to generate only a single, weak burst, with a strong decay of AP amplitude (not shown). This smaller drop observed in Ecel1 models is because they have a lower lever of I_T_ conductance in basal dendrites, which is negatively correlated with this feature. This mismatch with experimental data may be due to oversimplification of the T-type Ca^2+^ channel kinetics we model with a generic I_T_ conductance.

Even with some limitations, we could sample valid models for the three cell subtypes (Ecel1, Spp1 and Runaway) with a single run of MCMC, as opposed to relying on three separate sampling runs, each incorporating an adjusted mean of electrical features in the cost function to focus on a particular subtype. Therefore, we can study this ‘continuum of models’ to characterize their differences and similarities in terms of electrical features and parameters of intrinsic cellular mechanisms.

### Distinguishing electrical subtypes of models from model features and parameters

We begin by distinguishing electrical cell subtypes based on their electrical characteristics related to bursting properties. The first key metric is the burst number (see Method for its computations), which distribution shown in Fig. 3a is bimodal (see also Fig. 2f). We observe a peak at low burst numbers for Ecel1 models and at around 70 busts for runaway models. In the experimental recordings, because of a shorter recording time (14 seconds for experimental traces vs 25 seconds for MCMC) the second peak is at lower burst numbers.

**Figure 3:**
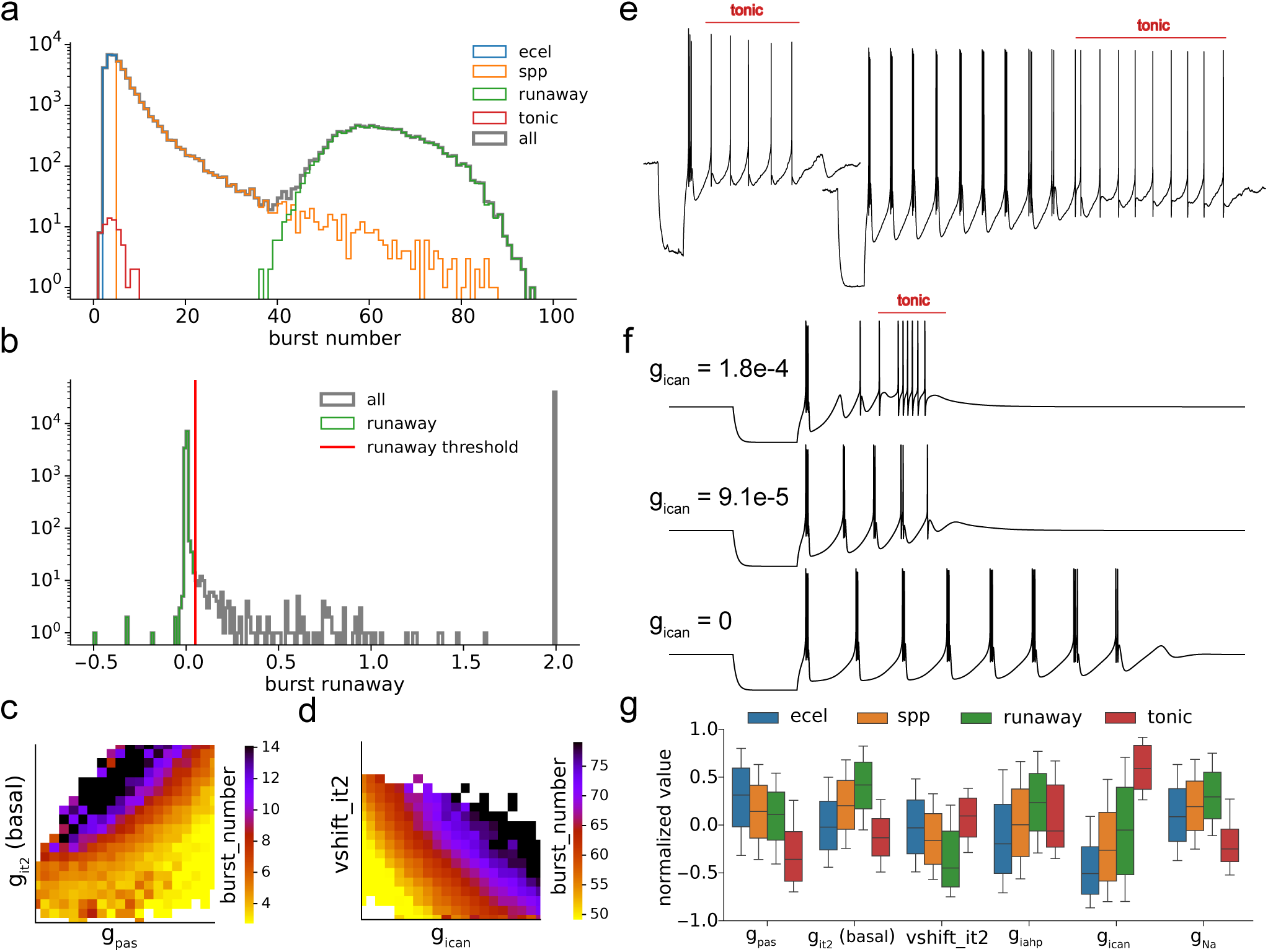
Electrical types analysis with the models. **a.** Distribution of burst number with three cell types and cells with tonic spiking after bursting. **b.** Distribution of runaway measure to detect runaway (smaller than 0.05). Values of 2 indicate large runaway values or undetectable ones. **c.** Average burst number corner plot of Ecel1+/Spp1+ burst number models with two mostly correlated parameters. **d.** Average burst number corner plot of burst number runaway models with two mostly correlated parameters. **f.** Example of experimental traces with transition to tonic (Ecel1+ left, Spp1+ right). **g.** Transition to tonic firing after burst in a model (top trace) and when I_CAN_ is decreased (lower traces). **g.** Distribution of parameters across cell types, for the statistically significant parameters only.

To distinguish Ecel1 from Spp1 models, we establish a threshold based on a burst number of 5, which corresponds to the experimental observations in the data (see Fig. 1d). It’s important to note that this classification is not strictly rooted in biological rationale, as there exists a recognized continuum of burst numbers among Ecel1+/Spp1+ cell types. Furthermore, some cells express both Ecel1 and Spp1 genes, positioning them along a transcriptomic gradient that spans the functional range between Ecel1 and Spp1 profiles, as detailed in Li et al. [2020]. Our sampled models similarly reveal this continuum, ranging from strictly Ecel1-like low-bursting to the Spp1-like high-bursting electrical type.

Then, to distinguish Runaway models from Spp1 models, a separate threshold of approximately 40 bursts could be implemented. However, this approach would incorrectly classify Spp1 models with high burst frequencies as runaway cells. We thus define an additional electrical feature, the burst runaway metric, that measures the slope of the slow AHP depth between bursts (see Fig. **??**b and Methods for exact feature definition).

We observe that traces with a burst runaway metric of less than 0.05*mV/ms* correspond to models characterized by bursts that remain stable over time and are unlikely to cease rebound bursting for an extended duration. This thresholding technique separates the second peak of models in the burst number distribution, exposing a long tail in the burst distribution for Spp1 models that exhibit high-frequency bursts (refer to Fig. 3a). We find that runaway cells generate action potentials towards the far end of the curve, as illustrated by the time to last spike feature summarized in SI Fig. 12a. Examining this parameter space allows us to obtain additional insights into the bimodal nature of the burst number distribution.

In Fig. 3c and d, we display the average burst number for a pair of the most highly correlated parameters associated with this feature, for the transitions from Ecel1 to Spp1 and from Spp1 to Runaway models, respectively. Each transition is governed by a unique set of correlated parameters to control this electrical feature. To elaborate further, for low bursting cells, the levels of I_T_ conductance and g pas play a crucial role in regulating the observed burst count. In contrast, for models firing a higher number of bursts, the conductance of I_CAN_ and the positioning of the T-type ‘window current’ (a phenomenon referring to the sustained influx of Ca^2+^ through Ca^2+^-gated ion channels, relying on the overlap between steady-state activation and inactivation [Williams et al., 1997]) become significant factors.

These differences stem from the fact that for runaway models, the burst number represents only the burst frequency, while for non-runaway ones the burst number also captures how fast the rebound bursting ends. Due to this qualitative difference, it is expected that a different set of parameters would control the burst number in these two regimes. In Fig. 3c, we observe larger burst counts for larger I_T_ conductance in the basal dendrites, Ca^2+^ influx is stronger and thus more rebound bursts are expected, while for larger passive conductance, the rheobase is also larger, hence Ca^2+^ influx is less likely to generate Na spikes in the soma. In Fig. 3d, a larger voltage shift (corresponding to an I_T_ current window centered at more negative voltages), results in a higher frequency of bursts because the rebound bursts are generated at a lower voltage and the membrane potential will take less time to get hyperpolarized after a burst to trigger the next one, see Cain and Snutch [2010] for a similar discussion.

A larger amount of I_CAN_ conductance also results in a higher number of bursts as I_CAN_ depolarises the cell once it is hyperpolarized after a burst, effectively shortening the time between two bursts. These trends will also be revisited in Fig. 4 in the context of burst curves, and in SI Fig. 12b where we show a prediction of an XGBoost regression model of burst number from model parameters. In these two regimes, we obtain a good accuracy (fraction of resp. 0.927 ± 0.002 / 0.956 ± 0.001 of correct prediction of burst number for resp. low/high groups) only when fitted separately from each other, confirming that the burst number feature captures different underlying mechanisms, depending on the cell type.

**Figure 4:**
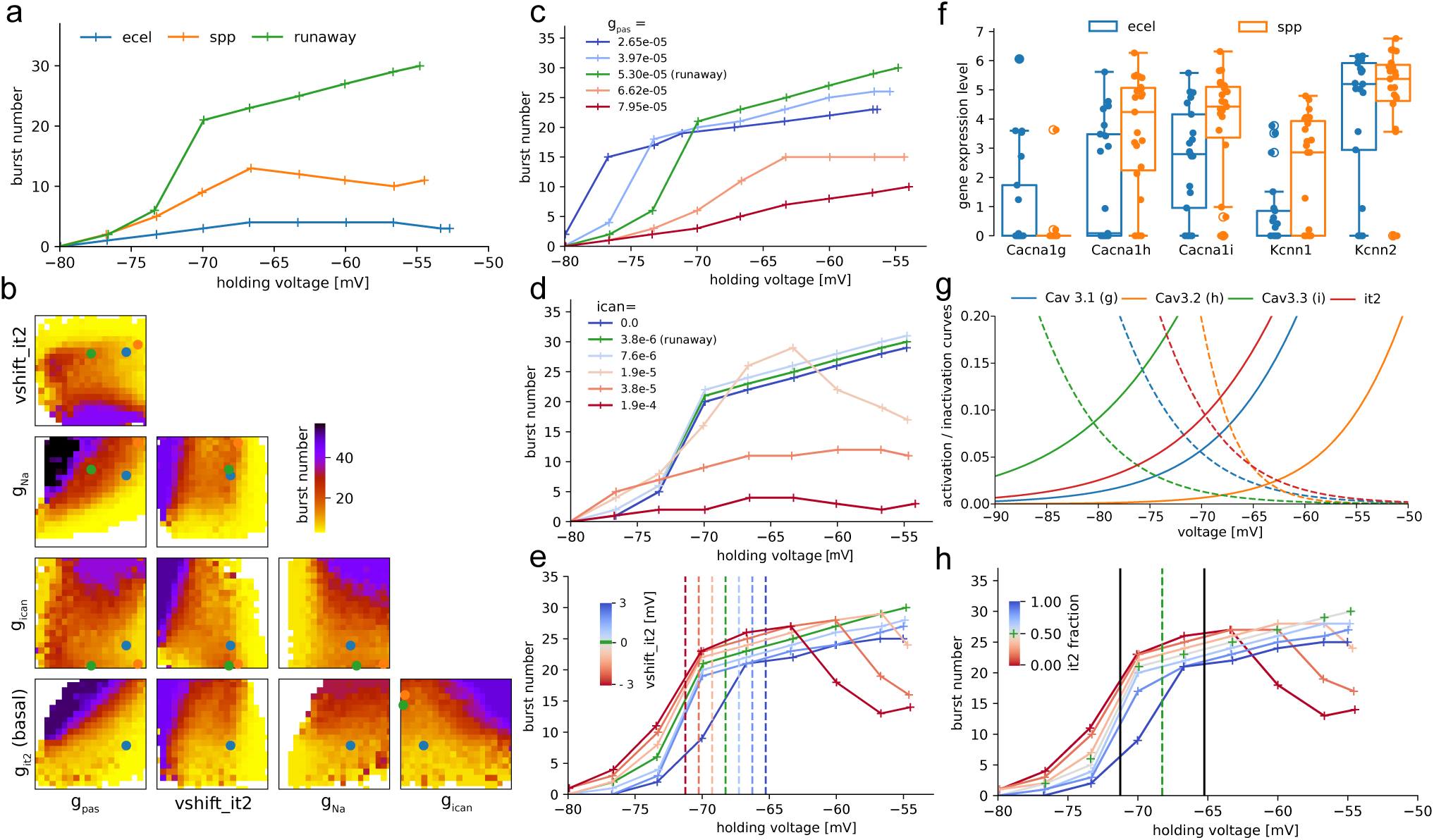
Burst number curve. **a.** Burst number curve for an Ecel1, Spp1 and a runaway model (different from the models of the previous figures). **b.** Corner plot of average burst number for most important parameters. Three models of previous panels are shown with dots. **c.** Change of g-pas affecting burst number curve of runaway model (green). **d.** Change of I_CAN_ affecting burst number curve of runaway model (green). **e.** Change of I_T_ voltage shift affecting the burst curve of runaway model (green). **f.** Gene expression levels (from Li et al. [2020]) of three genes related to Cav3 subtypes and two SK subtypes between Ecel1+ and Spp1+ cells. **g.** Activation and deactivation curves of subtype channels from McRory et al. [2001] and generic I_T_ used in this work. **h.** Replacing a single I_T_ with two I_T_ at 3mV shift left and right, with half conductance (gray line) is close to the model with a single generic I_T_ channel (green crosses). Relative fractions (colored lines) of each channel have a similar effect to a voltage shift (extreme curves are the same as in panel c).

Experimentally, it is common to see tonic spiking after rebound bursting, such as shown in Fig. 3e, but in our models only a small fraction display tonic action potentials after the initial rebound busting period. To study this behavior, we designed a specific feature to count the number of such tonic action potentials, and classify models with positive values as ‘tonic‘. This feature is delicate to define and misses cells with a low number of tonic AP, explaining part of our apparent under-sampling of these models, shown in red in Fig. 3a. One of the defining parameters for this behavior is a large value for I_CAN_ (see SI Fig. 12c,d). To assess this, we picked a tonic model in Fig. 3f (top) and reduced I_CAN_ to 0 (bottom) gradually, and observed the transition from an Ecel1 model with tonic, to an Spp1 model. Having enough I_CAN_ is not sufficient for the presence of tonic firing, it also needs low g pas, low I_T_ in basal dendrites, and low Na, as seen in Fig. 3g displaying the distributions of parameters between each cell type in the models. Generally, from Ecel1 to runaway models, we observe an increasing amount of I_T_, I_AHP_, Na and I_CAN_ conductances, with larger voltage shifts and lower g pas values, confirming our previous analysis. The voltage shift difference between cell types is because, for larger voltage shifts, a cell will have a smaller burst number, as a smaller fraction of T-type Ca^2+^ channels are open at the holding voltage of −65*mV* (see Fig. 4g). These cells will therefore be more often classified as Ecel1, from this protocol with holding voltage at −65*mV*.

### Burst curve variability and T-type Ca^2+^ channels

We now study the relationship between burst curves and T-type Ca^2+^ channels, particularly burst curve variability induced by the relative fraction of conductances of the Cav3 channel subtypes. First, we show in Fig. 4a the burst curves of three electrical models of each subtype, selected from those with the maximum burst number at intermediate voltages. We show the burst number for a simulation time of 14s, as in the experimental data. For this duration of recordings, most data points above 15 bursts are of runaway type (with a small runaway metric). In the experimental data summarized in Fig. 1c-e, we observe a large variability in the shape of burst curves, in particular in terms of the location and amplitude of its peak. We aimed to determine if our models, which incorporate only the generic T-type Ca^2+^ conductance and omit the specific T-type Ca^2+^ channel isoforms expressed in reticular neurons in vivo, could accurately represent these characteristics of the burst curve. First, in Fig. 4b we present a corner plot of the average number of bursts across pairs of parameters that mostly control this feature. These correlations emerge for traces at a fixed holding voltage but will also nonlinearly control the shape of the entire burst curve (i.e., the number of observed bursts per holding voltage).

While we cannot examine all of these correlations in detail, we will focus on two of them more thoroughly. First, in Fig. 4c, we consider the effect that modifying g-pas has on the burst curve. The effect of g-pas on burst number exhibits a nonlinear relationship, with a decrease in the number of observed bursts at both high and low values of g-pas. For high values (orange/red curve), the runaway regime is lost, while for low values, the width of the peak is so large that the cell rebound bursts for the entire trace at a holding voltage of −55mV (all points above 15 bursts are runaway).

This is compatible with the previous result presented in Fig. 3c, where large g pas is correlated with a low number of bursts. The passive conductance of the membrane, denoted by g pas, represents how easily ions can flow across the neuronal membrane through passive channels, impacting the membrane potential and its response to inputs. In the lower range of g pas values (blue/green traces), increasing g pas corresponds to faster dynamics when the cell is hyperpolarized (between bursts) resulting in shorter intervals between rebound bursts and larger burst numbers. With a large enough increase of g pas, the hyperpolarization period becomes too short to deactivate enough Ca^2+^ channels to then trigger a strong enough low threshold spike and subsequent rebound burst (red curves), resulting in lower burst numbers. Second, we plot in Fig. 4d a similar effect upon altering I_CAN_, also seen earlier in Fig. 3d. In this case, I_CAN_ depolarizes the cell, hence also reducing the time interval between bursts for low values while preventing rebound bursting for larger values. While the trends are similar with g pas, the mechanisms are different, g pas speeds up the dynamics, while I_CAN_ pushes the membrane potential to higher voltages.

Another crucial property of the I_T_ conductance for the burst curve is the location of its window current, or the range of voltages at which there is a significant likelihood of Ca^2+^ influx through the channels, effectively creating a window for current to flow. In Fig. 4e we show the impact of varied voltage shifts parameter of the I_T_ channel on burst number (green is the base runaway model). Apart from an expected shift of the burst curve, the maximum number of bursts (or burst frequency) also changes (see Fig. 3d). For larger voltage shifts, which result in the window current being adjusted to lower voltages, the burst curve also shifts and reaches its peak at these lower voltages. For smaller voltage shifts, there is not a sharp decrease in burst numbers below −55mV but an overall reduction of the burst frequency where the runaway type of steady bursting starts at higher holding voltages.

Relying solely on traces obtained from a single holding voltage leads to degenerate models since numerous burst curves can have the same burst number at that given voltage. As a result, to create a model of a specific cell, one would need to consider the entire burst curve and remove this degeneracy. Here, we are rather interested in capturing the variability of the experimental burst curves, hence we only use three holding voltages, with large standard deviations of target burst numbers. This variability in the burst curves observed among cells of the same type indicates that these cells must possess different relative fractions of conductances to replicate the experimental phenotypes. So far, our model only included the generic T-type Ca^2+^ channel conductance, but with a voltage shift to account for some of the variability of the burst curve. Voltage shifts of ion channel models are often fixed in agreement with experimental conditions (such as liquid junction potential) at which these channels have been recorded. In our models, this voltage shift accounts for the relative conductance of the T-type Ca^2+^ family or Cav3 channel subtypes, as we will demonstrate below.

A single-cell transcriptomics analysis of Ecel1+ and Spp1+ neurons reported differences in gene expression related to voltage-gated ion channels known to be important shapers of neuronal rebound burst firing ([Li et al., 2020]). Ecel1+ cells exhibited higher expression of the Cacna1g gene, which encodes the low-voltage-activated T-type calcium channel subunit *α*-1g. In contrast, Spp1+ cells had increased expression of the Cacna1h and Cacna1i genes, encoding the *α*-1h and *α*-1i isoforms, respectively (summarized in Fig. 4f).

Direct electrophysiological comparison of *α*-1g, *α*-1h, and *α*-1i under identical recording conditions and identified unique functional characteristics of these three isoforms, suggesting distinct contributions to neuronal physiology [McRory et al., 2001]. Examining the window current for each channel isoform illustrated in Fig. 4g, we see it peaking at progressively more hyperpolarized voltages for *α*-1h, *α*-1g, *α*-1i. Although all three isoforms overlap in their steady-state activation and inactivation properties, those relative differences in the expression level could produce the observed differences in the burst curve characteristics of Ecel1+ and Spp1+ cells. This relation between the relative fraction of Cav3 channel subtypes and the location of the burst curve is visible in Fig. 1e, where the center of the burst curve is at higher voltages for Ecel1+, consistent with the higher expression of Cacna1g gene (Cav3.1 channel subtype) for this cell subpopulation. There were also differences in the expression of the small conductance Ca^2+^-activated K+ (SK) channels, which are responsible for the afterhyperpolarization (AHP) phase of rebound burst firing. Spp1+ TRN neurons expressed both SK1 and SK2 channel variants, whereas Ecel1+ neurons predominantly expressed the SK2 subtype [Li et al., 2020] (see SI Fig. 15 for more gene expression comparison between cell types). Due to differences in the activation and inactivation kinetics, as well as Ca^2+^ sensitivity of these SK channel isoforms, TRN neuron subpopulations exhibit varying AHP time scales based on their specific SK channel isoform expression profiles.

As a first approximation, we treat the Cav3 channel subtypes as voltage-shifted versions of the generic I_T_ conductance, disregarding any variations in characteristics like time constants. By substituting the single generic I_T_ with two copies that are shifted up and down by the same voltage, *v*_shift_, and halving the overall maximal conductance, we develop a cell model comprising two Cav3 subtypes rather than just one generic channel. For small enough *v*_shift_, that is when the window currents of the shifted channel still overlap significantly, the artificial subtype model should remain similar to the single channel model. In the limit of vanishing *v*_shift_, it will converge to the generic model. In Fig. 4h we conducted this experiment with *v*_shift_ = 3*mV* and indeed, the burst curves remain similar. Next, by introducing an imbalance in the conductances of the up and down-shifted I_T_ model, as illustrated in the same panel, we can manipulate the position of the burst curve, similar to how we adjusted the voltage shift of the single channel in Fig. 4e. This numerical experiment shows that the voltage shift of the generic I_T_ conductance can be considered an approximation of the relative conductances of the Cav3 channel subtypes. This approach could be used to relate the differential contribution of ion channels to single-cell dynamics in the context of pathologies. For instance, the upregulation of Cav3.2, associated with the absence seizures in epilepsy [Cain et al., 2018], would have the primary effect of increasing the amplitude of the burst curve around −65*mV* (Fig. 4g), due to an increase in the magnitude of the overall T-type Ca current around that voltage range (Fig. 4a), explaining the hyperexcitability and enhanced burst firing that underlie absence seizures. Dysregulated Cav3 channel activity is also associated with spindle and slow-wave impairments in schizophrenia, with a localized reduction in Cav3.3 activity in TRN as one potential cause of observed spindle abnormalities [Thankachan et al., 2019]. Such a targeted reduction of Cav3.3 currents could be tested in Ecel1 or Spp1 model neurons in isolation, to assess its impact on the burst firing properties of different TRN neuron subtypes.

### Model generalizations to biological temperature and morphological population

After having reproduced the biological variability of rebound burst firing in TRN with our models, we turn to the study of their generalization to higher temperatures (from 25 to 34 Celsius) and a population of detailed reconstructions for the same morphological type. This generalization will enable us to use these models in a thalamoreticular microcircuit at physiological temperature, with morphological variability from Iavarone et al. [2023] and study the resulting network dynamics in the next section.

First, in Fig. 5a, we investigate the effect of temperature in our models. We constrained the models at 25 degrees Celsius, the temperature used in experimental recordings, but we intend to use them at 34 degrees in the circuit simulation since this is the physiological temperature. To implement this adjustment, we utilize the q10 temperature coefficient integrated into each ionic mechanism model (see Method). As illustrated in Fig. 5a (insets), the primary impact of raising the temperature from 25 to 34 degrees is a reduction in the burst count, accompanied by an increase in both the frequency and the number of action potentials within each burst, along with a decreased time to the first spike. Due to the absence of experimental recordings and their electrical characteristics at 34 degrees, we relied on the temperature dependence to develop a baseline model capable of replicating the observed burst numbers for each cell type.

**Figure 5:**
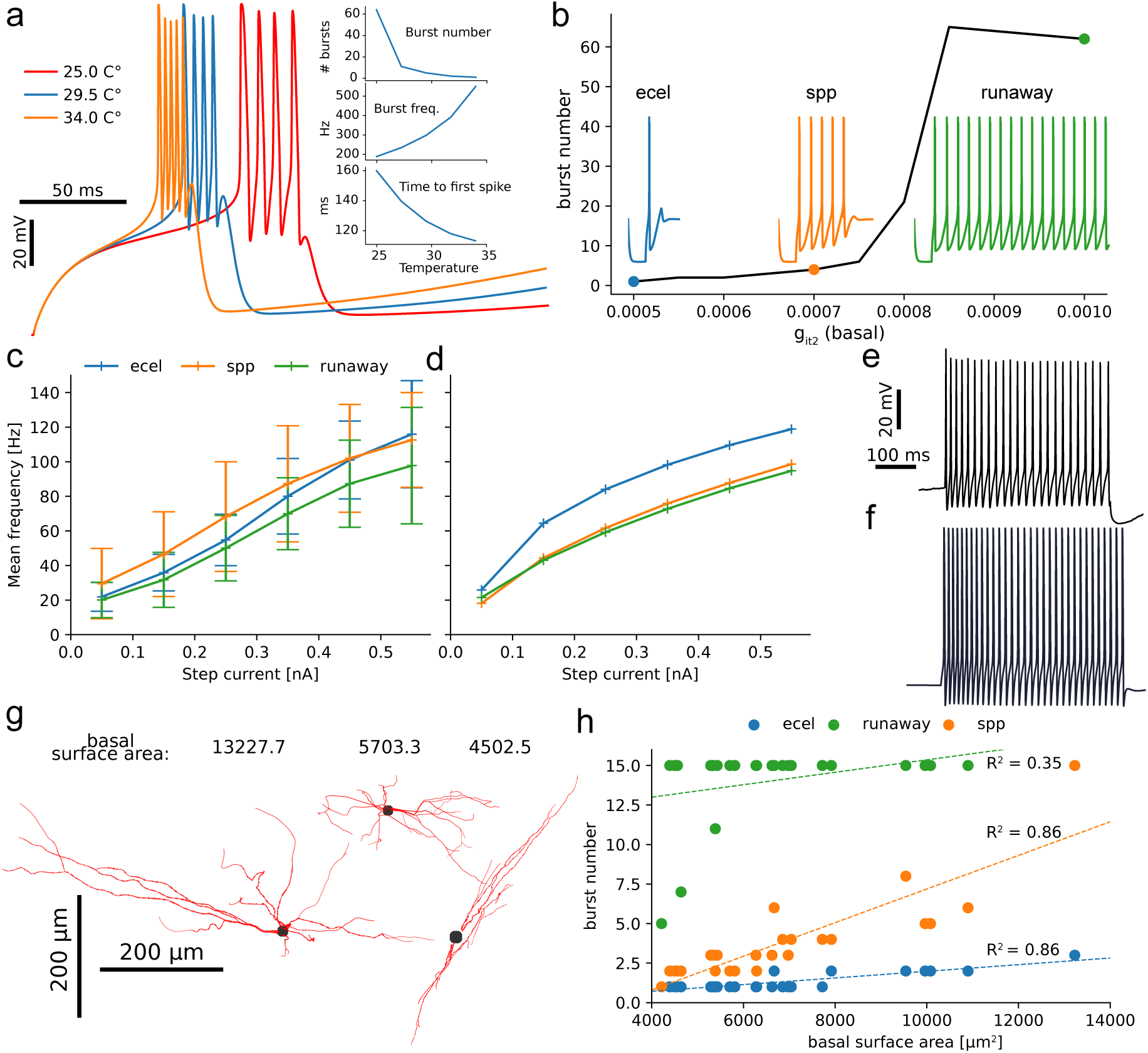
Generalization of electrical models for circuit simulations. **a.** Traces of bursts when increasing the simulation temperature from 24 to 34 degrees Celsius. Insets are three features (burst number, burst frequency and time to first spike as a function of temperature, computed on the full traces (not shown). **b.** Number of bursts for a fixed model with the I_T_ conductance in basal dendrites varied. Three models are highlighted which will be used as Ecel1/Spp1 and runaway representatives. Traces show rebound bursting of each of the three models. **c.** Experimental IF curves for all three types during tonic protocol at 25 degrees Celsius **d.** IF curve of tonic firing for the three models shown in Fig. 2 at 25 degrees Celsius. **e** Representative experimental trace of tonic regime for an Ecel1+ cell under a 0.15nA step current. **f** Representative model trace for the Ecel1 model under a 0.15*nA* step current. **g.** Example of detailed morphological reconstructions [Iavarone et al., 2019] with various total dendritic surface areas. **h.** Burst number for the three models on all reconstructions as a function of their dendritic surface area. The burst number is clipped at 15, and the dashed lines are linear regression with associated *R*^2^ values.

Starting from a model with a low score (below 2 std) at 25 degrees from the MCMC sampling, we created a baseline model at 34 degrees (see column circuit model in Table 2). To compensate for the loss of burst number due to higher temperature (see Fig. 5a top inset), we mainly increased the I_T_ channel conductance (see also Table. 2. Varying the I_T_ conductance in the basal dendrites of this baseline model covers the full range of observed burst numbers of all subtypes (black curve in Fig. 5b). We then selected three I_T_ values along this curve to obtain three baseline models, one for each subtype. Simply altering the I_T_ conductance among the subtypes is an oversimplification; however, it enables us to consistently examine the impact of I_T_ conductance in circuit simulations by minimizing variability from other parameters.

In our model development using the MCMC sampling approach, we explicitly excluded any tonic spiking protocols, even though this firing mode plays a crucial role in the network-level activity. Therefore next, we validate our models by comparing the current frequency (IF) curve at 25 degrees Celsius with the experimental IF curves. This comparison is illustrated in Fig. 5c, where the data has been binned and averaged across all cells of the same type. We do not notice any substantial differences in IF curves across cell types, however, Ecel1+ cells exhibit a greater tendency to enter a state known as depolarization block when subjected to high current injections. We did not attempt to model this phenomenon here, see Hartley et al. [2024] for further details on the experimental data. In Fig. 5d, we plot the IF curves for the three models at 25 degrees presented in Fig. 2, demonstrating a strong consistency with the experimental data presented in the previous panel. Experimental traces were maintained at −60mV before the step current injection, so we replicated this condition in the model to ensure consistency.

In our model, we observe that Ecel1 has a higher firing frequency since our random choice of models resulted in a near-maximal Na conductance in the soma while the Spp1 and Runaway models have near minimal values of Na in the soma (see Fig. 2c and Table 2). The IF curve of these three models therefore bounds the range of possible IF curves of the models, in agreement with the variability in the experimental data (see Fig. 5c). This indicates that the relationships between parameters and features can be utilized to modify electrical models as needed, specifically somatic Na and excitability in this context.

Fig. 5e-f shows representative tonic firing traces for both experiments and models, elicited by a 0.15*nA* current injection. Both exhibit similar action potential shapes and firing frequencies. Our models, constructed without explicit matching of experimental IF curves, nonetheless reproduce them. This suggests that IF curves and neuronal excitability are inherent properties of the burst protocol, rather than additional constraints.

We then investigated the role of underlying morphology in determining the validity of each electrical type. In Fig. 5g we show three examples of basal dendrites from our dataset of 25 morphologies [Iavarone et al., 2023]. Some have a smaller dendritic surface area compared to others, yet exhibit a similar level of branching complexity (see SI Fig. 9). In Fig. 5h, we present the burst numbers for our three electrical models when assessed across the population of morphologies, plotted against their dendritic surface area. For each cell type, dendritic surface area correlates significantly with burst number, while soma surface area shows a weaker, but still noticeable, correlation. (not shown, slopes and r2 of *s* = 0.00087 *r*2 = 0.5 for Ecel1, *s* = 0.0037 *r*2 = 0.42 for Spp1 and *s* = 0.00141 *r*2 = 0.17 for Runaway). Therefore, with few exceptions of extremely small or large morphologies, cell types retain their distinct burst number characteristics across the entire morphological dataset. Additionally, the observed correlation between burst number and I_T_ conductance sensitivity suggests that dendritic I_T_ is more critical for rebound bursting than somatic I_T_, consistent with prior research [Crandall et al., 2010].

### The contribution of Ecel1, Spp1 and Runway model neurons to spindle-like oscillations

Thalamic spindle oscillations, frequently observed during cortical up states [Destexhe Alain, 2007, McCormick and Bal, 1997, Steriade M, 1993], arise from a mechanism involving TRN-mediated inhibition of thalamic relay cells [Iavarone et al., 2023]. To delve deeper into the role of specific TRN cell subtypes, we utilize the thalamoreticular microcircuit connectome (Fig. 6a) and examine the differential impact of Ecel1 and Spp1 TRN cells on spindle properties.

**Figure 6:**
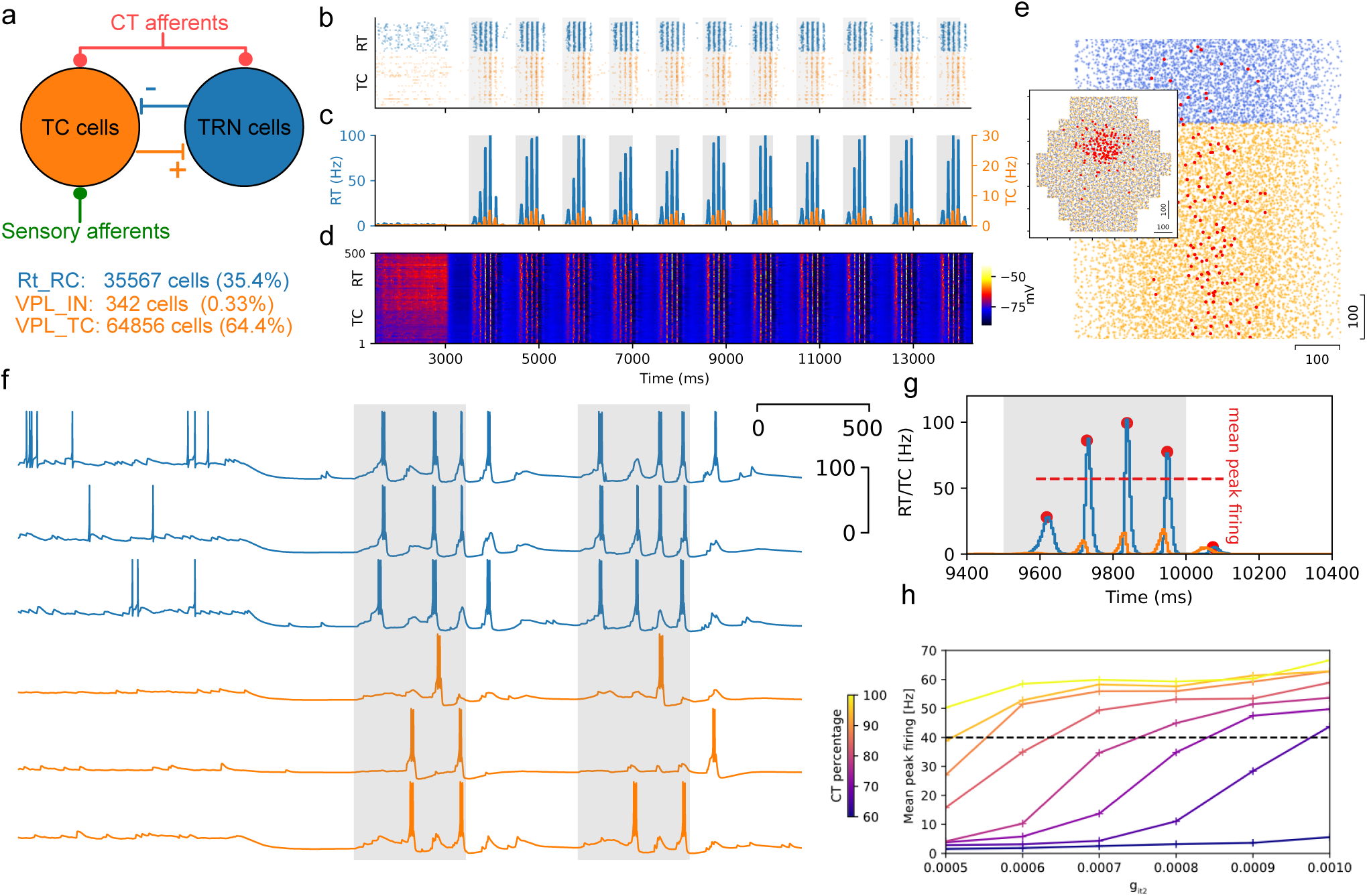
Simulations of spindles. **a.** Sketch of the circuit connectivity and composition. **b.** Spike raster plot (500 neurons), gray shadows represent up states. **c.** Peri-event time histograms (PETH) (500 neurons), gray shadows represent up states. **d.** Voltage raster plot (500 neurons). **e.** Spatial distribution of cells in the microcircuit column (blue for RT, orange for TC cells), efferent cells of a single CT afferent are highlighted in red (top view of the column in inset). **f.** Voltage traces of three sample cells per electrical type during the awake state and the two first spindles. **g.** Zoom on one spindle, showing the AP frequencies of the two cell types (a zoom on panel b). Red dots are ripple peaks, with average mean peak firing values. **h.** Spindle mean peak firing across various circuits with uniform models in a range of I_T_ conductances, for different fractions of CT inputs.

To achieve this, we replaced the original reticular models [Iavarone et al., 2019] with varying proportions of Ecel1, Spp1, and runaway models. The original model further subdivided reticular cells into adapting and non-adapting types, but as their contribution to spindle behavior was deemed less critical, we focused on non-adapting cell types only. We have created three circuit variants, replacing all TRN cell models with Ecel1, Spp1 and/or Runaway model neurons to test these circuits’ ability to produce spindle-like oscillations in a simulated NREM-like state. Fig. 6b-d presents simulation results for a sample of TRN and TC cells from a circuit where all TRN cells are modeled as Ecel1 neurons. In this simulation, we observe spindle-like oscillations brought by simulated cortical up states, reproducing the results of Iavarone et al. [2023] (see Fig. 4C). In our study, we extended the sequence of simulated cortical up and down states to 16 cycles, as a reasonable trade-off between simulation duration and accuracy of averaged quantities such as mean peak firing. This measure is computed as the average firing rate of the peri-event time histograms (PETH) of each spindle event, summarized in Fig. 6g. The average and standard deviations of the mean spindle peak firing during a simulation are further used to characterize the ability of a circuit to produce spindle-like activity following the onset of simulated slow oscillations in cortical afferents.

In the model of Iavarone et al. [2023], all the placed CT synapses present in the circuit are recruited to participate in cortical up states, input which we refer to as 100% CT activation. In this study, we were interested in characterizing how differences in intrinsic cellular excitability of TRN subpopulations contribute to emergent network dynamics. To this end, we varied the strength of synaptic depolarization by recruiting different percentages of CT afferents during each up state, starting from 40% up to 100%. Fig. 6h presents the mean peak firing levels for several intensities of CT synaptic inputs and the range of dendritic I_T_ conductances. The electrical models depicted in Fig. 5b were reused, with only the dendritic I_T_ conductance being modified to span the full range of the three cell types, from an Ecel1 model to runaway models with maximal I_T_. For weak CT activation below 60 − 70%, peak firing levels are minimal, indicating an inability of the circuit to reliably generate spindle-like oscillations at this level of synaptically driven depolarization. Stronger CT activation results in more pronounced peak firing levels, particularly in circuits with higher I_T_ conductance levels. At 100% CT input, all circuits exhibit similar peak firing levels above 40Hz (dashed line), a threshold indicative of large spindle-like oscillations. This result demonstrates a strong correlation between the amount of cortical synaptic input received by the thalamus, the overall Ca^2+^ currents in TRN cells and the ability of the circuit to generate spindles.

### I_T_ and I_AHP_ conductances predict the mean peak firing rate of spindle-like oscillations

The TRN is segregated into modality-specific sectors based on its innervation of particular thalamic nuclei and reciprocal connections with specific cortical areas [Pinault, 2004]. This anatomical organization results in regionally specific oscillatory properties, which could account for differences in local sleep-wave activity. It has been demonstrated that oscillatory burst firing varies across TRN sectors, with sensory sectors exhibiting more repetitive burst firing compared to the limbic sector [Fernandez et al., 2018]. Hence, one critical aspect we address with this modeling work is the impact of the TRN circuit composition in terms of Ecel1/Spp1 ratio, as well as the presence of runaway cells to the spindle dynamics. Building on the previous section’s uniform TRN circuits, we created mixed circuits with varying cell compositions to explore the relationship between effective I_T_ conductance and mean peak firing during spindle-like oscillations.

Fig. 7a-b shows circuit activity with increasing Ecel1 fractions at 90% CT input. Higher Ecel1 fractions, with lower total I_T_ conductance, lead to decreased peak firing. While peak firing decreases, high-firing spindles still occur, albeit less reliably, as seen in the variability for high Ecel1 fractions (Fig. 7c).

**Figure 7:**
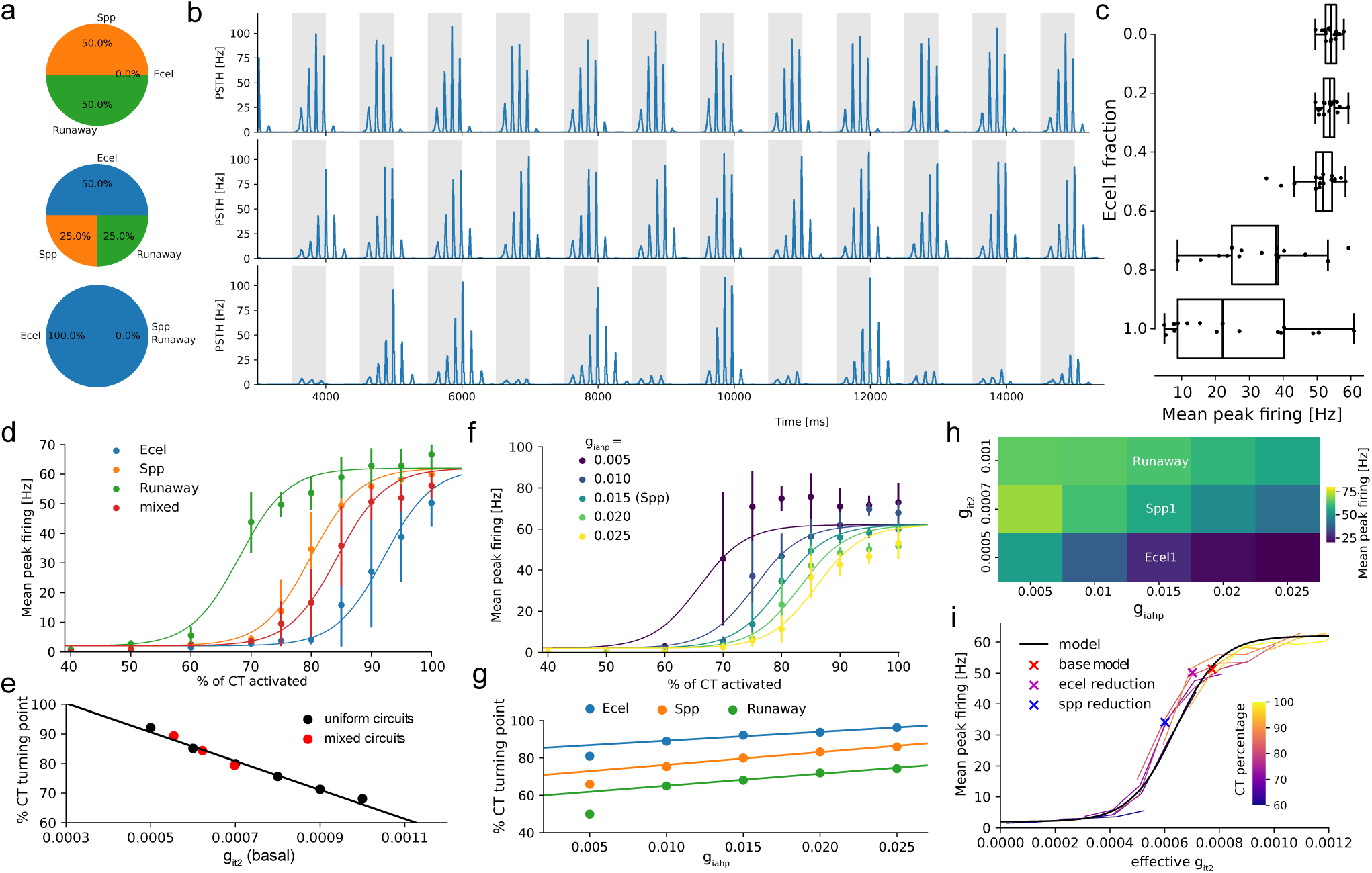
Peak firing of spindle-like oscillations can be predicted from circuit composition. **a.** Illustration of three circuits with different cell compositions, varying the fraction of Ecel1 cells.fix cakes with overlapping **b.** TRN spikes PSTH for each of the three circuit variants in panel a during cortical up/down states (highlighted in gray). **c.** Spindle peak firing as a function of the fraction of Ecel1 cells in six circuits. Each dot is a spindle during a simulation (16 per simulation). **d.** Spindle peak firing as a function of CT input for three uniform circuits with the three base models, and a circuit with Ecel1 fraction of 0.5 from panel c. We fit only the position of a sigmoid function (CT turning point), all other parameters are kept the same. **e.** Sigmoid centre from panel d, or CT turning point as a function of effective circuit I_T_ conductance (weighted harmonic mean of models basal dendrite I_T_ conductances). The black line is a linear fit. **f.** Spindle peak firing as a function of CT activation levels for circuits with various levels of I_AHP_ conductance. **g.** Same as panel e but with varying I_AHP_ conductances for each of the three base models. Linear fit ignores the point at 0.005 as bursts are not clearly defined in the simulation. **h.** Average spindle peak firing as a function of I_T_ and I_AHP_ conductances in uniform circuits at 90% CT activated. add ecel/spp/runaway to y axis, mark iahp value of these models **i.** The black line represents the predicted peak firing at 85% from the sigmoid fit and position from panel g. Coloured lines are simulation data from Fig. 6h, shifted along the x-axis by using the slope of panel g. Three crosses are simulations using mixed models, reproducing the experiment of Li et al. [2020]Fig. 5.

To gain deeper insights into the relationship between the level of I_T_ conductances in basal dendrites and mean peak firing during spindle-like oscillations, we plot in Fig. 7d the mean peak firing levels as a function of CT synaptic input for three uniform circuits (Ecel1, Spp1 and runaway models) and one mixed circuit model composed of half Ecel1 and half Spp1/runaway models, as illustrated in the second row of panels Fig. 7a-b. For each of these circuits, the dependence of peak firing on CT input exhibits a sigmoidal relationship, with the primary variation occurring in the location of the turning point, which is influenced by the level of I_T_ conductance in the respective model neurons. To quantify this relationship, we fitted sigmoid functions to the data from 6 uniform models, fixing all parameters except for the turning point location. The resulting turning point values were then plotted against the corresponding dendritic I_T_ conductance values in Fig. 7e. This relation is linear, with high confidence, hence the amount of I_T_ currents in basal dendrites is predictive of the mean peak firing observed during spindle activity in these circuits. To further analyze the mixed circuits depicted in Fig. 7a-b, we applied the same sigmoid fitting procedure and computed the effective I_T_ conductance for each cell model. The effective I_T_ conductance was calculated as the weighted harmonic mean of the dendritic I_T_ conductances of the individual models, weighted by their relative fractions within the circuit. The use of the harmonic mean is appropriate given the density-like nature of conductances. As illustrated in Fig. 7e (red dots), effective I_T_ conductance emerges as a reliable predictor of the mean peak firing during spindle-like oscillations, following the same linear trend as observed for circuits with uniform cell models (black dots). This finding suggests that the level of peak firing during spindle activity can be predicted from the effective I_T_ basal dendrite conductances across various circuit compositions.

Next, we characterize the effect of varying the conductance of SK channels, which are responsible for the afterhyperpolarization (AHP) current, on peak firing. When activated, SK channels hyperpolarize reticular cells, enabling them to repeatedly rebound burst fire while controlling their burst frequency and regularity [Cueni et al., 2008]. In the TRN, downregulation of SK channels has been shown to increase burst duration and frequency, whereas upregulation increases AHP amplitude and reduces intrinsic excitability, as demonstrated by Silvan et al. [2024] and SI Fig. 14. In Fig. 7f-g, we present a similar analysis of the observed mean peak firing while varying the level of I_AHP_ conductances in basal dendrites. We observe similar sigmoid profiles, although with an opposite trend. For a fixed level of CT synaptic input, increasing the level of I_AHP_ conductance results in a decrease in peak firing.

At the lowest level of I_AHP_ conductance (at 0.005), the circuit does not exhibit regular and well-defined spindle-like oscillations. Instead, the neurons burst for most of the simulation duration, irrespective of the level of cortical inputs. This behavior can be attributed to the shorter hyperpolarization periods associated with low I_AHP_ levels, which allow the cells to trigger subsequent bursts more rapidly. As the level of I_AHP_ conductance increases, each burst comprises fewer action potentials, and the subsequent hyperpolarization period becomes longer, making it less likely for the the cell to burst on subsequent ripples of the spindle (Fig. 14d, f). This results in reduced average peak firing for the same level of CT input (see Fig. 7f-g). In Fig. 7g we perform a linear fit (excluding the lowest I_AHP_ point at 0.005) for the CT turning point of I_AHP_ scan of Fig. 7f, similar to the fit in Fig. 7e. We conducted the same analysis for the other two uniform-composition circuits, composed of either Ecel1 or Runaway cells, and observed a similar positive slope. This demonstrates that I_T_ and I_AHP_ conductance levels have opposite effects on the mean spindle peak firing and can be used in concert to predict the mean peak firing in a circuit during spindle-like activity (see Fig. 7h for a map of the spindle peak firing at 90% CT input). The linear dependence between the sigmoid turning point and the level of I_T_ conductance (see Fig. 7e) allows us to establish a mapping between cortical synaptic input and effective I_T_ conductance for maintaining a constant spindle mean peak firing frequency. This enables us to replot the data from Fig. 6h, showing I_T_ scans at various CT input fractions, as a function of this effective I_T_ conductance. This replotting validates our prediction of spindle peak firing frequency, as the curves superimpose onto the sigmoid model at 85% of CT input, as depicted in Fig. 7i.

This predictive power of the effective I_T_ conductance enables us to constrain the cell compositions of a circuit from its mean spindle peak firing, or rather spindle robustness or reliability. For example, it was found that a reduction of Ca^2+^ currents in Ecel1+ cells has only a marginal effect on the average number of spindles, but a similar reduction of Ca^2+^ currents in Spp1+ cells induces a significant decrease in the number of spindles [Li et al., 2020]. In Fig. 7i (crosses), we show the result of a similar experiment in our model. We built a circuit with 10% Ecel1 cells (red cross), then reduced I_T_ conductance by 0.0002 independently in Ecel1 (magenta cross) and Spp1/runaway (blue cross) to create two additional circuit variants. Only the circuit with the reduction in Spp1 cells has an effective reduction of I_T_ large enough to decrease the mean spindle peak firing, similar to Li et al. [2020]Fig. 5. Hence this modelled correlation between I_T_ conductances and spindle peak firing may serve as an implicit constraint on TRN sector cell composition from experimental data. In this case, the constraint is on the maximal fraction of Ecel1 cells of about 10 − 15% such that reducing I_T_ conductances in Spp1 cells produces a larger drop in effective I_T_ than the reduction in Ecel1, and reproduces the result of Li et al. [2020].

### Conclusion and outlook

In Li et al. [2020], and more recently in Hartley et al. [2024], neurons in the thalamic reticular nucleus have been experimentally studied in the context of understanding how intrinsic cellular properties impact emergent network-level dynamics. It was found that reticular neurons can be segregated into two transcriptionally defined populations, Ecel1+ and Spp1+, named after two genes most deferentially expressed by these neuronal subpopulations. Forth of these experimental results, we tasked ourselves with creating detailed numerical models of these cell types and investigating their properties, from ion channels to circuit dynamics with spindle-like oscillations. Leveraging the sampling method for building electrical models of detailed neurons [Arnaudon et al., 2023], we obtained a large pool of models matching the experimental variability of the electrophysiological features across the two cell types. Consistent with the findings of Li et al. [2020], we did not observe a clear-cut distinction between Ecel1+ and Spp1+ TRN cell types, but rather a continuum of models with a smooth transition in properties, such as maximal rebound burst firing. Furthermore, the electrophysiology dataset underlying our modeling effort contains a significant proportion of cells that exhibit prolonged bursting activity, referred to as runaway neurons [Hartley et al., 2024]. We were able to generate models of these cells by sampling from the same MCMC parameter space, and by exploring it, we gained insights into the ion channels that primarily control the transitions between different cell subtypes. While T-type Ca^2+^ and SK (I_AHP_) channels are the most important ion channels in shaping the bursting behavior of TRN neurons, the balance between active and passive membrane properties, such as the ratio of passive to active conductances (g_pas_ /I_CAN_ or g_pas_ /g_Na_), also plays a crucial role. These passive properties can influence the overall excitability of the neuron and, consequently, its ability to generate bursts.

Due to a lack of complete and coherent ion channel models that account for the different isoforms of T-type Ca^2+^ and SK channels, we were constrained to using more generic models. To address this limitation, we considered a strategy involving duplicated generic I_T_ channels, which were symmetrically shifted in voltage. We demonstrated that the relative conductances of these duplicated channels correspond to a voltage shift of a single generic I_T_ channel. This result indicates that, as a first approximation, the diverse properties of different channel subtypes can be modeled by adjusting the density and voltage sensitivity of a generic channel model. By manipulating these parameters, we can effectively simulate the behavior of different channel isoforms. This simplified approach proved particularly useful in capturing a significant portion of the variability observed in the burst curves, which quantify a cell’s ability to rebound burst when subjected to hyperpolarizing current injections. Building on the work of Iavarone et al. [2023], where a microcircuit of the somatosensory thalamus was built and validated to subserve spindle-like activity, we have incorporated Ecel1 and Spp1 electrical models into the TRN tear of the circuit and asked how intrinsic ionic conductances modulate network activity. We found that both I_T_ and I_AHP_ conductances of the TRN cells are accurate predictors of peak firing during spindle-like oscillations. Each has an opposite effect on the spindle peak frequencies and balances each other to maintain a certain level of bursting. For circuits composed of multiple cell subtypes (mixed circuits), an effective I_T_ conductance (defined as the harmonic weight mean of I_T_ conductances) is the measure predictive of spindle peak firing. It would be of interest to also study mixed circuit variants with different levels of I_AHP_, which we predict to follow a similar, but opposite trend.

This allowed us to reproduce the knock-out experimental result of Li et al. [2020]Fig5 whereby the reduction of Ca^2+^ currents in Ecel1+ cells had a non-significant effect on spindle oscillations but a similar modification in the Spp1+ subpopulation visibly reduced spindle numbers, spindle duration, and led to fragmented sleep. Reproducing this experimental result also gave us upper bounds on the cell composition of the circuit, and in particular on the maximum number of Ecel1 cells as compared to Spp1 cells of around 10 − 15% such that this effect could be observed. This bound in cell composition cannot be directly compared with experimental data but may serve as a basis for predicting the cellular composition of different TRN tiers based on localized sleep-related patterns in modality-specific thalamocortical loops [Fernandez et al., 2018].

Furthermore, such a numerical model can be used to test hypotheses linking abnormalities in sleep spindles to alterations in ion channels within specific TRN subpopulations, which may underlie the pathophysiology of a number of disorders. Several sleep studies report marked deficits in sleep spindles as well as slow-wave abnormalities in schizophrenia [Castelnovo et al., 2018, Zhang Y, 2019, Thankachan et al., 2019]. Interestingly, in our model, we observe that circuits composed of an increasingly higher fraction of Ecel1 neurons, intermediate strength CT up states don’t provide sufficient synaptic excitation to reliably initiate a spindle on every up state, without failures. It would be instrumental to investigate in more detail the mechanisms behind these “borderline” regimes, where spindles are generated less robustly. From a translational perspective, developmental disruptions in the expression of risk genes, particularly those highly expressed in the TRN and associated with disorders like schizophrenia (e.g., Cav3.3), could potentially alter the balance of Ecel1-like and Spp1-like neuron, impacting spindle generation through broader transcriptional changes. This raises important questions about the influence of ion channel mutations, such as those in Cav3.3, on the developmental trajectory of these neuronal subtypes. While Cav3.3 loss-of-function might shift neuronal phenotypes towards an “Ecel1-like” bursting mode, it could also significantly impact cell composition and the overall architecture of thalamic circuitry. A deeper understanding of these mechanisms may pave the way for novel biomarkers and targeted therapies for these disorders.

While we have been successful in creating accurate detailed single-cell models and reproducing network-level dynamics observed experimentally, our models rely on several assumptions that could be refined with additional experimental data. First, the use of generic T-type Ca^2+^ and SK channels limited our ability to precisely capture isoform-specific properties, particularly in reproducing highly skewed burst curves. Other kinetic factors, such as variations in time constants, may be crucial for accurately modeling these complex firing patterns. Second, while we incorporated a linear increase in the density of the I_T_ conductance in the basal dendrites, we did not conduct a detailed analysis of the specific impact of this distribution on the electrical models. Integrating experimental data from dendritic recordings [Crandall et al., 2010] could help address this limitation and assess the impact of non-uniform conductance distributions on cellular electrical properties. Such an approach could be further linked to morphological properties of cellular subtypes, as some studies report different ratios of surface areas between somatic and dendritic compartments [Harding-Jackson et al., 2022]. Furthermore, we did not attempt to reproduce the depolarization block state of Ecel1+ cells, which was frequently observed at high current amplitudes during tonic firing in the experimental data [Hartley et al., 2024]. As the implications of this depolarization block for circuit-level function are not well-characterized, we leave this aspect for future investigations, which will likely require more detailed models of Ca^2+^ /SK and Na+ channels. Additionally, the adaptation of data from 24 to 34 degrees Celsius, while practical, introduces assumptions that warrant further experimental validation. Lastly, the findings presented here are specific to the somatosensory TRN and may not generalize to other sectors.

While we generated a diverse population of single-cell models through MCMC sampling, our initial focus was on a select few models to gain a clear understanding of the relationship between specific ion channels and spindle properties. To fully capture the complexity of TRN network dynamics, incorporating a diverse population of single-cell models with varying properties, as suggested by [Arnaudon et al., 2023], is essential. This approach would allow to further explore how individual cell-to-cell variability impacts network-level behavior. Overall, we believe our work provides a solid foundation for future numerical studies of detailed reticular models, from ion channel dynamics to network-level properties, enabling a deeper understanding of the underlying mechanisms of thalamic function.

## Acknowledgment

This study was supported by funding to the Blue Brain Project, a research center of the École polytechnique fédérale de Lausanne (EPFL), from the Swiss government’s ETH Board of the Swiss Federal Institutes of Technology.

## Methods

### Electrophysiologcal data

The experimental data is from Hartley et al. [2024]. In short, neurons were subjected to 500ms long hyperpolarizing current injections at various holding potentials ranging from −80 to −50 mV, with the negative current adjusted to hyperpolarize each cell to a range of −110 to −100 mV.

### Morphological reconstructions

The morphological reconstructions used in this work for single cell and circuit simulation are the same as in Iavarone et al. [2023]. For Rt cells, they consist of 25 detailed reconstructions cloned into 7315 morphologies for building the microcircuit. These morphologies were reconstructed and processed by Iavarone et al. [2023], including unravelling and repair procedures to account for the various biases induced by the reconstruction techniques.

To perform MCMC sampling of electrical models, we have to select a representative morphology, and we did so by following the approach of Arnaudon et al. [2023], as an average morphological model from the population of detailed reconstruction, see SI Fig. 9. We first created a model for the soma with a single cylinder of average radius and length so the surface area matches the average from the population (see SI Fig. 9a-b). We then estimated the average surface area density of the basal dendrites as a function of the path distance to the soma from all the reconstructions and selected the morphology closest to this average (green in SI Fig. 9d and Fig. 2b). We noticed that soma and dendritic surface areas are correlated in this dataset (*R*^2^ = 0.6), with some variability in the areas, number of neurites and maximum branch orders, see SI Fig. 9e.

### Burst curves fitting

The burst number of each trace is computed using the feature described below and the holding voltage is estimated as the average voltage during the first step of the bursting protocol. We fit this data with a skewed normal distribution

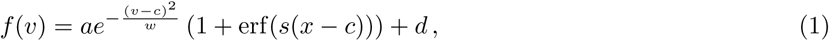

with skewness parametrized by *s*, width by *w*, amplitude by *a*, centre by *c* and linear shift by *d*. We use the following bounds for these parameters in the fitting function (Scipy curve fit function): *a* ∈ [0, 100], *w* ∈ [20, 180], *c* ∈ [−100, −40], *s* ∈ [−2, 2] and *d* ∈ [0, 1].

### Classification of cell types

From the extracted electrical features of recording during bursting protocols, we applied the XGBoost classifier with SHAP feature importance analysis Lundberg and Lee [2017] to detect which electrophysiological feature was most important to distinguish electrical types. Doing a 10-fold cross-validation of traces with maximal burst number per cell using the entire set of electrical features extracted from rebound burst protocol data yielded an accuracy of 88.2 + −13.7%. Excluding the most obvious differentiator, maximum burst number per holding membrane voltage, and reclassifying the traces using the rest of the electrical features still produced similar accuracy results (86.0 + −15.7%). Finally, using only the top three features identified as most influential, namely burst number, burst mean frequency and spike width allowed us to classify the traces with an accuracy of 87.6 + −14.3%. The top features in each classification with their Shapely values are shown in SI Fig. 10.

### Ion channel models

For the building of detailed electrical models, we assigned to the somatic and bacal compartments the following list of mechanisms.

1. hh2 Na and hh2 K from Destexhe et al. [1994] (soma). Fast Na and K channels to make Na spikes, *ena* = 20*mV* and *ek* = −80*mV* were fixed to approximately match AHP depth and AP amplitude from data
2. cad from Destexhe et al. [1994] (soma and basal): Ca^2+^ dynamics with linear decay term, decay rate fixed to 150ms for both to approximately match the shape of slow AHP.
3. I_CAN_ from Destexhe et al. [1994] (basal): builds up to stop bursting and transition to tonic firing
4. I_T_ from Destexhe et al. [1994] (soma, basal with increasing density away from soma): generic T-type channel that creates low threshold Ca^2+^spikes, has an additional parameter for the slope of linear increase. It has a voltage shift parameter, where positive values shift towards negative voltages.
5. I_AHP_ from Destexhe et al. [1994] (soma and basal): or SK channel, responsible for the slow hyperpolarisation after bursts, voltage shift of 10mV to the reversal potential was applied to match the depth of slow AHP from data
6. I_A_ from Huguenard and McCormick [1992] (basal): stops bursting early, not studied in detail in this work
7. g pas (all): passive channels with *e pas* = −70mV. For I_T_, the increasing density function is

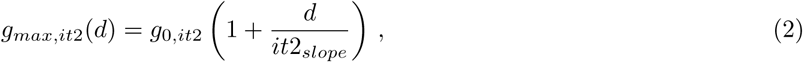

where *d* is the path distance from the soma.

### Protocols and features

The protocol for simulating rebound bursting is as close as possible to the experimental protocol. First, for a period of 5 seconds, we inject a current to maintain the cell at the specific holding voltage. This current was previously found by a bisection search based on a step protocol of 2 seconds long. With the same bisection search, we find the current to hold the cell at −100*mV* and use it for a 500*ms* hyperpolarisation step. After this step, we keep the initial holding current for 25 seconds. For MCMC sampling, we used three holding voltages, −80*mV* to ensure no bursts are created, −65*mV* as the main protocol around the peak of the burst curve, and −55*mV* to prob the burst curve around the largest possible voltage.

From these protocols, we compute the following features: the burst number (see below for details on its computation), number of spikes per burst, burst mean frequency (frequency of spikes in each burst, averaged over all bursts), peak voltage (absolute voltage of the peak), AP2 AP1 peak difference (the drop of spike amplitude between the first two spikes), time to first and last spike, first ISI (inter-spike interval, other ISIs are highly correlated with this one), AHP depth (in absolute voltage, averaged across all spikes), postburst minimum values (or slow AHP depth between bursts, averaged across all bursts), voltage std (std of voltage trace to ensure it is stable), tonic after burst (number of tonic spikes after the bursting period, see below) and three Ca^2+^ concentration amplitudes along the basal dendrite. In SI Table 1 we show the mean and std of these features that were used for MCMC sampling, set by hand, close to the experimental data.

### Burst number feature

The burst number is the number of bursts occurring during a trace. The definition of a burst from a trace is not trivial, especially for edge cases, such as for traces with a single burst, or bursts of single action potentials. We tried to design a feature robust to these edge cases as follows, see SI Fig. 11 for an illustration. We compute the inter-spike intervals between all AP of the trace and look for a clear bimodal distribution, distinguishing the inter- and intra-burst ISIs. The number of inter-burst ISIs plus one gives the number of bursts. For edge cases, if all ISIs are smaller than 50ms, we assume it is a single burst. If only a few bursts have more than one spike, it will be sufficient to detect the bimodal distribution, hence all the subsequent bursts of a single spike will be counted as bursts. If all the bursts have a single spike, it will not be counted as bursting, which is a rare edge case.

### Burst runaway feature

To detect if a cell is of type burst runaway, we use the time to last spike feature, but also a measure of the slope of voltage drift of slow AHP depth between bursts, see SI Fig. 11 for an illustration. We compute the slow AHP depth between each burst (discarding any possible minima before 5ms after the last AP in a burst), and compute the slope of the voltage drift between the second and the one before the last slow AHP depth. We discard the first and the last one, to prevent bias from some unusual start of bursting, cut traces at the end, or unusual transition to the end of the bursting period. If the drift is present, assumed to be if the slope is larger than 0.05mv/ms, it indicates that the bursting behaviour is not sustainable, and it will stop at some point (maybe after the end of the recording).

### Tonic number feature

Some traces stop bursting, but still fire action potential in a more regular, tonic manner. We have designed a feature able to detect these spikes not as bursts, but as tonic spikes, see SI Fig. 11 for an illustration. This is also used in the burst number feature, to ensure consistency. The number of tonic AP in a trace is detected as follows. In the bursting regime, the slow AHP depth between bursts is always lower than the fast AHP depth between APs, while for the tonic regime, they are of comparable depth. Hence, we count the number of detected inter-burst ISIs which have slow AHP higher than fast AHP depth. This gives us a good estimate of the number of tonic spikes in a trace (adding one to the number of tonic ISIs).

## Supplementary information

### A Supplementary tables

**Table 1:**
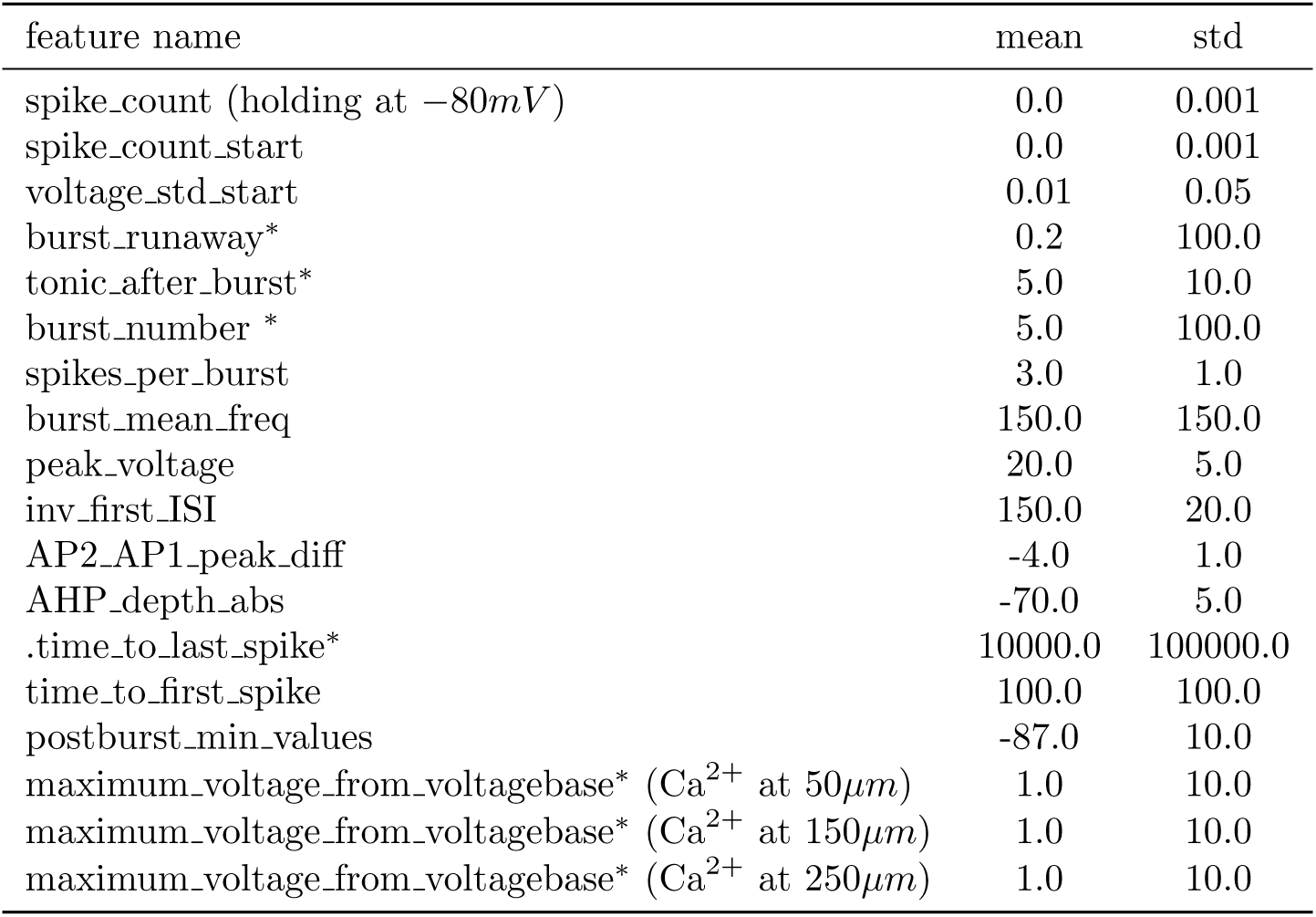
Electrical features names (as in eFEL, see https://github.com/BlueBrain/eFEL) and values mean and standard deviations. Features with a ^∗^ have a large std and can be considered as not constraining the MCMC sampling (as it used the largest score as cost).

**Table 2:** Parameter values of models used in the main text. The models at 25*C* are from MCMC, and the ones at 34*C* are for the circuit simulations (only the I_T_ in basal dendrites differs between them).

| Mechanism name | Added to | Ecel1 | Spp1 | Runaway | Ecel1/Spp1/Runaway |
| --- | --- | --- | --- | --- | --- |
| Temperature (T) | - | 25 | 25 | 25 | 34 |
| Passive conductance (g_pas) | soma/basal | 7.58e-05 | 4.194e-05 | 3.523e-05 | 3.0e-05 |
| Voltage shift of $I_T$ (vshift_it2) | soma/basal | -1.69 | -4.13 | -5.88 | -3.0 |
| Fast transient Na (hh2_Na) | soma | 2.88 | 1.17 | 1.5 | 2.0 |
| Delayed K (hh2_K) | soma | 0.76 | 1.28 | 1.60 | 1.0 |
| Low-threshold $Ca^{2+}$ ($I_T$) | soma | 0.00092 | 0.00088 | 0.00058 | 0.0005 |
| Slope low threshold $Ca^{2+}$ density (it2_slope) | basal | 517.16 | 357.01 | 418.0 | 300.0 |
| $Ca^{2+}$ -dependent K ($I_{AHP}$) | soma | 0.0701 | 0.0324 | 0.0285 | 0.03 |
| Slow $Ca^{2+}$ dependent nonspecific cation ($I_{CAN}$) | basal | 6.031e-05 | 9.689e-05 | 10.02e-05 | 4.0e-05 |
| $Ca^{2+}$ -dependent K ($I_{AHP}$) | basal | 0.000736 | 0.0138 | 0.0149 | 0.015 |
| Low-threshold $Ca^{2+}$ ($I_T$) | basal | 0.000667 | 0.00077 | 0.00072 | 0.0005/0.0007/0.001 |
| Transient voltage-gated K ($I_A$) | basal | 0.00144 | 0.0043 | 0.00195 | 0.00 |

### B Supplementary figures

**Figure 8:**
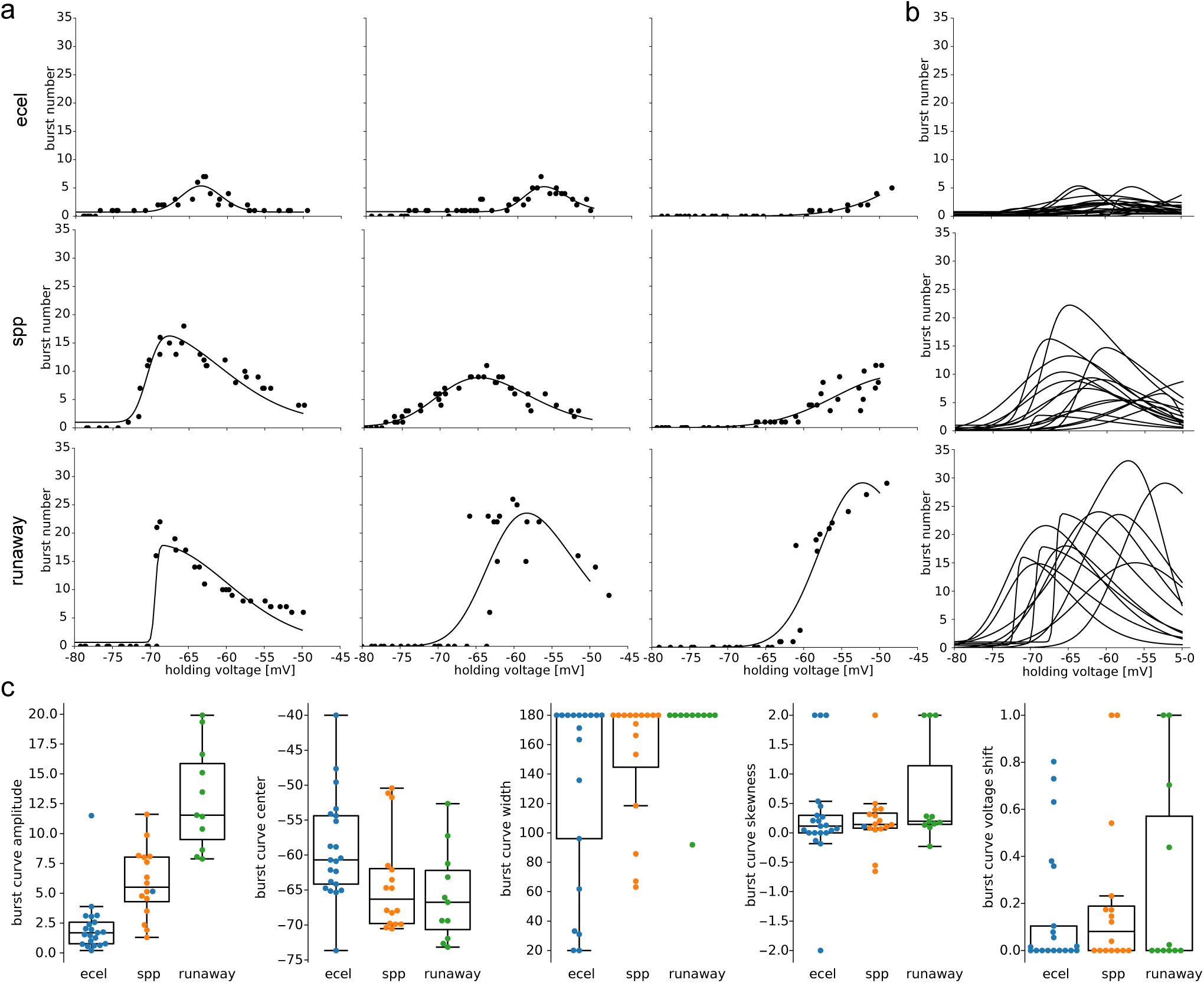
Variability in experimental burst curves. **a.** Three examples of experimental burst curves per cell type, illustrating cases with large skewness, or centred at large or low holding voltages. **b.** Superimposed fits of all burst curves, showing the large variability per cell type. **c.** Distribution of all the fit parameters of the burst curves, per cell type.

**Figure 9:**
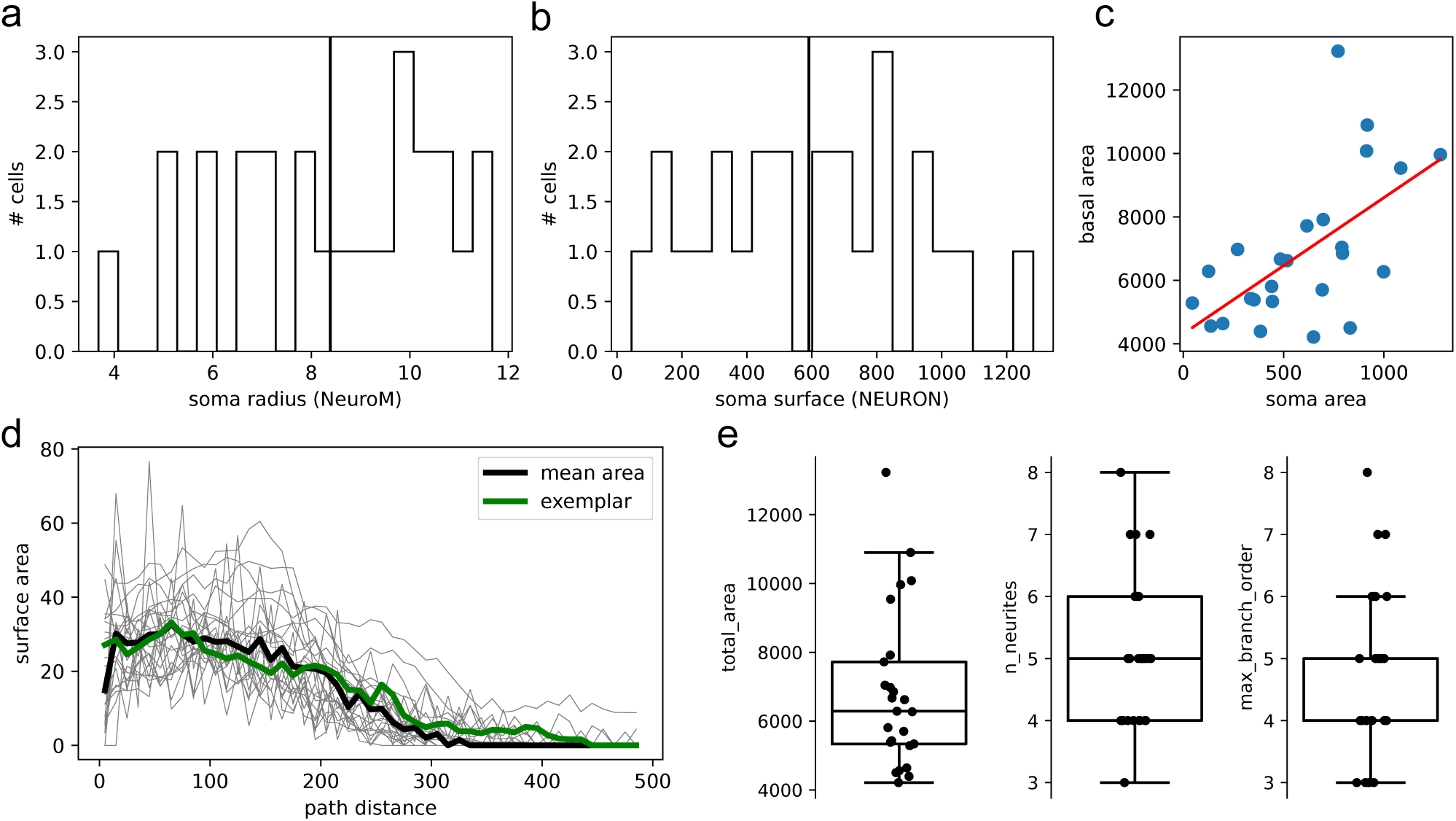
Exemplar morphology. **a.** Distribution of soma radii from the population of morphologies of Iavarone et al. [2023] **b.** Distribution of soma surface area from the same population. **c.** Surface are profiles of each morphology, the average of them, and chosen exemplar, closest to this average. **d.** Correlation between some surface area and basal surface area (person = 0.6).

**Figure 10:**
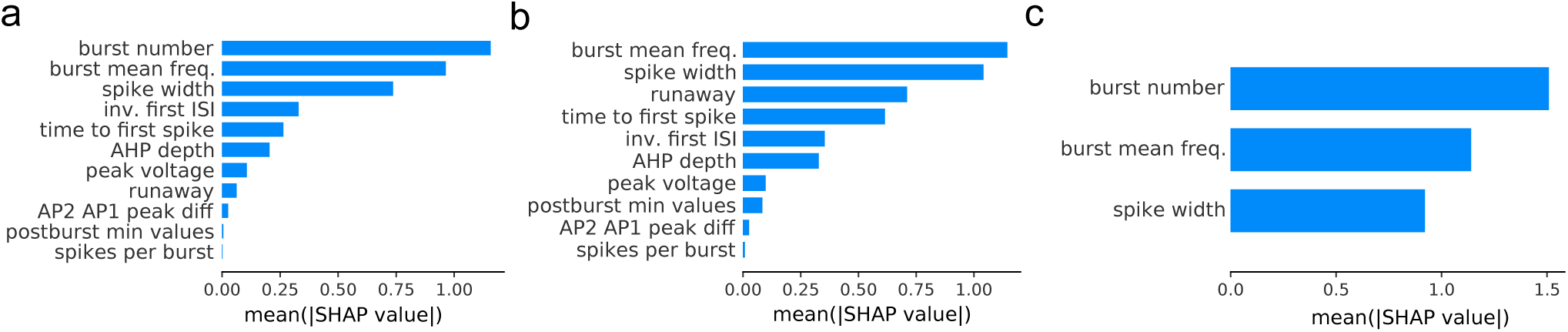
Feature importance on cell type classifications. **a.** SHAP values for a 10-fold classification using all electrical features extracted from the rebound burst protocol voltage traces (accuracy of 88.2 ± 13.7%) **b.** SHAP values for a 10-fold classification using electrical features as in a., but with maximum burst number per holding membrane voltage feature excluded (86.0 ± 15.7%) **c.** SHAP values for a 10-fold classification using the top three most influential electrical features (87.6 ± 13.8%).

**Figure 11:**
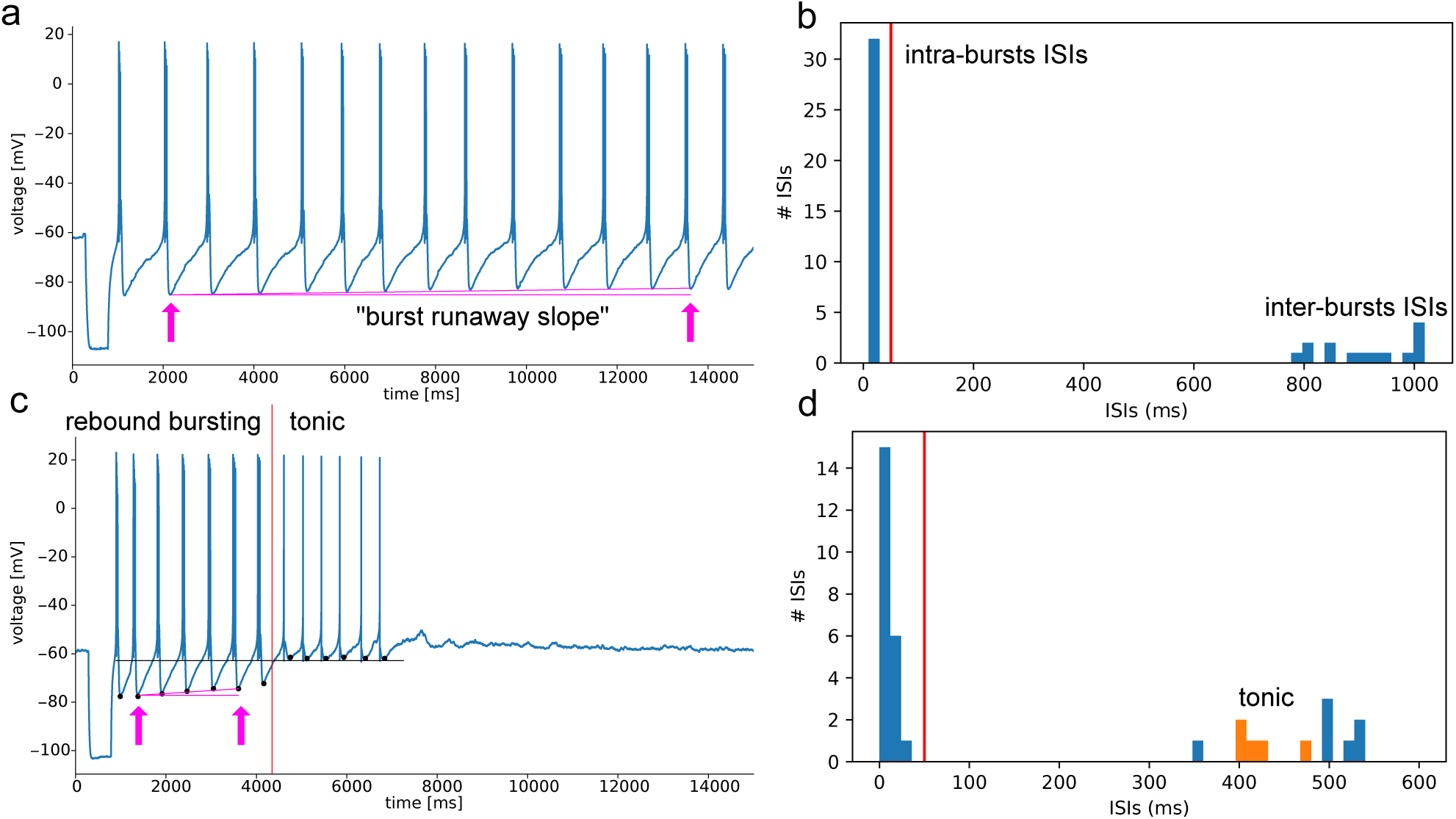
Electrical features for rebound burst characterisation. **a.** Rebound burst trace with first and last burst used to compute the runaway slope marked with arrows. The slope is computed from the slow AHP depths between these two bursts, assuming linearity. **b.** Inter-spike interval distributions of the trace in panel a, showing the small ISI within the bursts, and larger ones between bursts, defining the number of bursts (minus one). **c.** Trace of a cell rebound bursting, then transitioning to tonic spiking. Our runaway slope computation is also highlighted in panel a. The black dots show the slow AHP depths between bursts, and the black line the fast AHP depths between action potentials in the bursts. For bursts with slow AHP depths higher than this mean fast AHP depth, the burst will be considered tonic APs. **d.** From the split of bursts and tonic AP in panel c, we can differentiate the ISIs between burst and tonic AP and estimate the number of burst and tonic APs.

**Figure 12:**
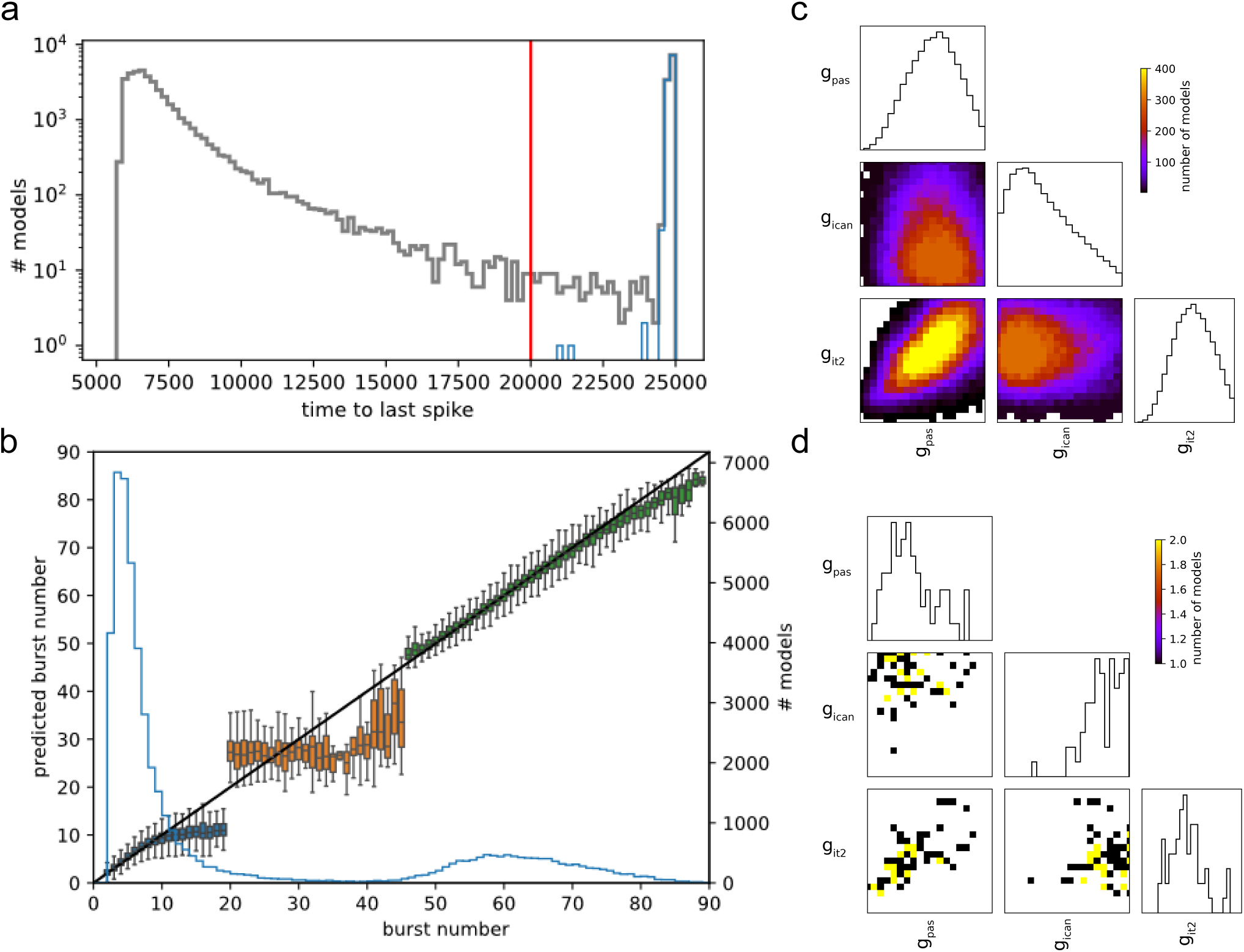
Additional panels for fig 2. **a.** Distribution of time to last spike with models in blue as runaway. **b.** Predicted burst number of an XGBoost regression in three parts, for low, middle and high burst numbers. A single model fails to fit the entire range, but low and high range work. **c.** Sub-corner plot of three main parameters for tonic firing after burst, for all models **d.** Sub-corner plot of same parameters for only models showing tonic after burst.

**Figure 13:**
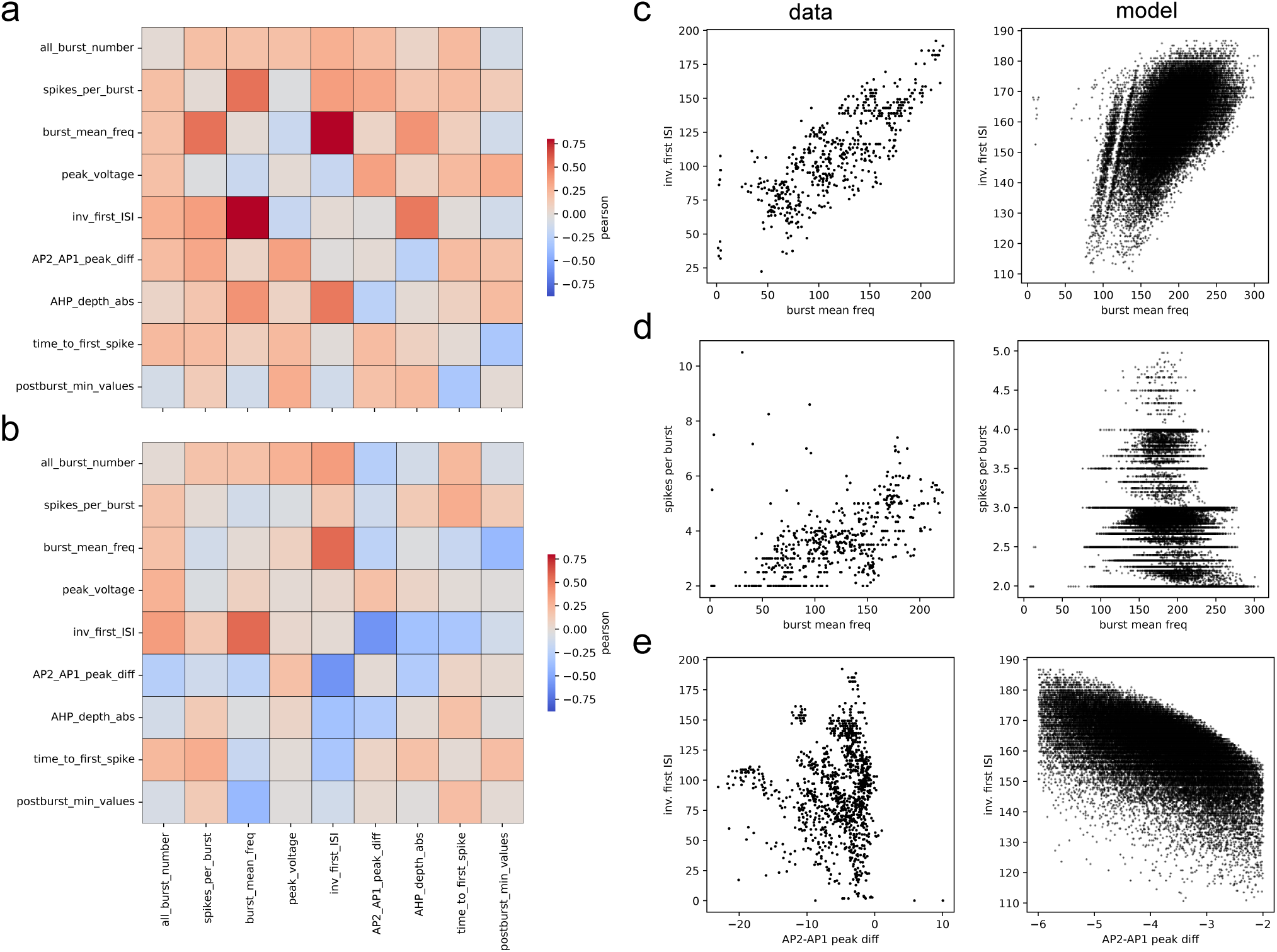
Two dimensional correlations between features in experimental data and models. **a.** Pearson correlations of experimental data **b.** Pearson correlations of MCMC models **c-e.** Scatter plot of some strong correlations in data (left panels) and models (right panels).

**Figure 14:**
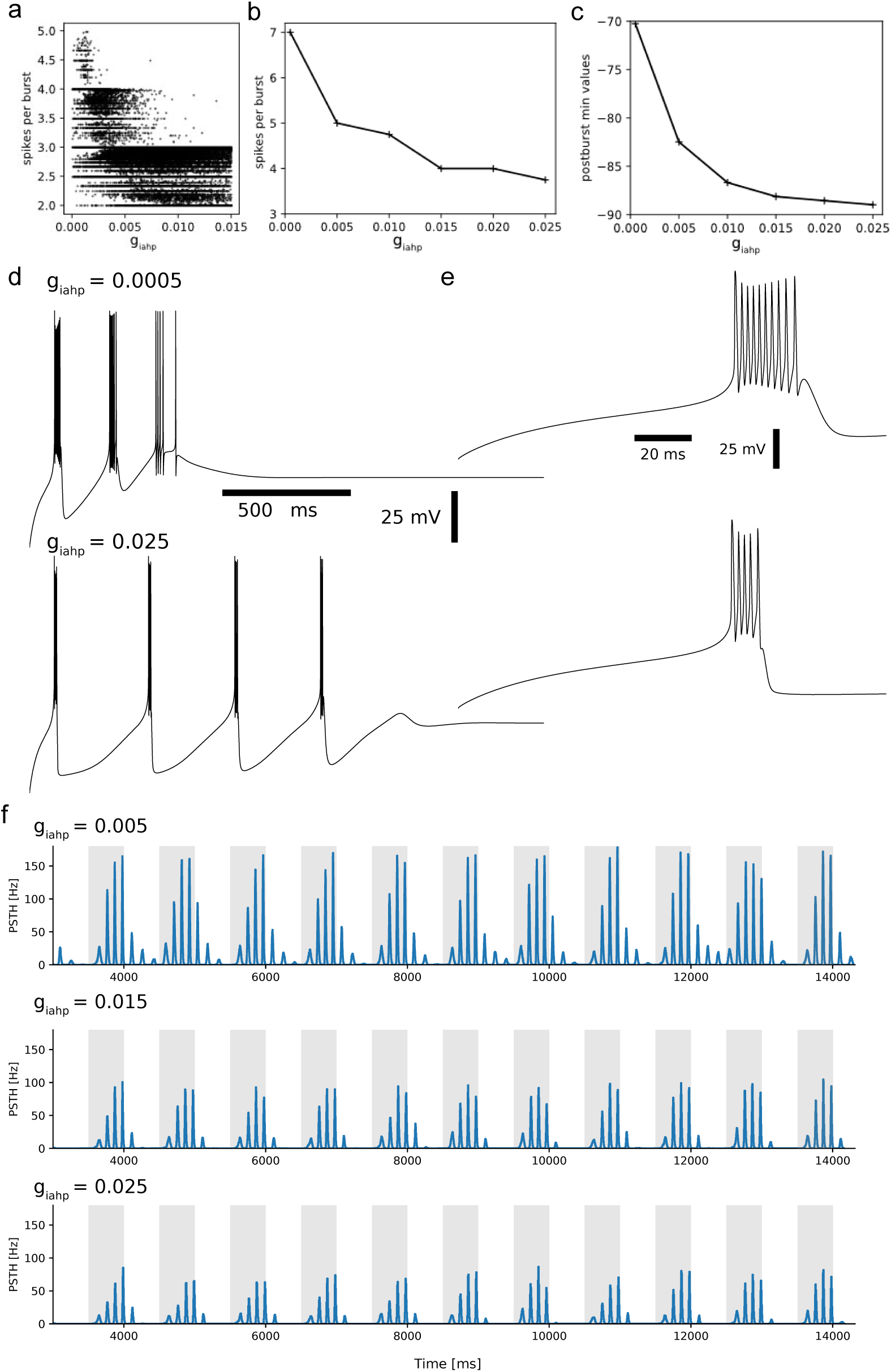
Effect of I_AHP_ channel. **a.** Correlation between I_AHP_ channel and number of AP per bursts in all MCMC models with cost *<* 2 std. **b.** Spike per burst as a function of I_AHP_ in the Spp model. **c.** Average Slow AHP depth between bursts as a function of I_AHP_. **d.** Traces for the model with lowest and highest I_AHP_ values from the previous panel. **e.** Zoom on the first burst of traces in panel d. **f.** Spindle simulations with three I_AHP_ conductances for the Spp1 uniform circuit (middle row is the Spp1 model).

**Figure 15:**
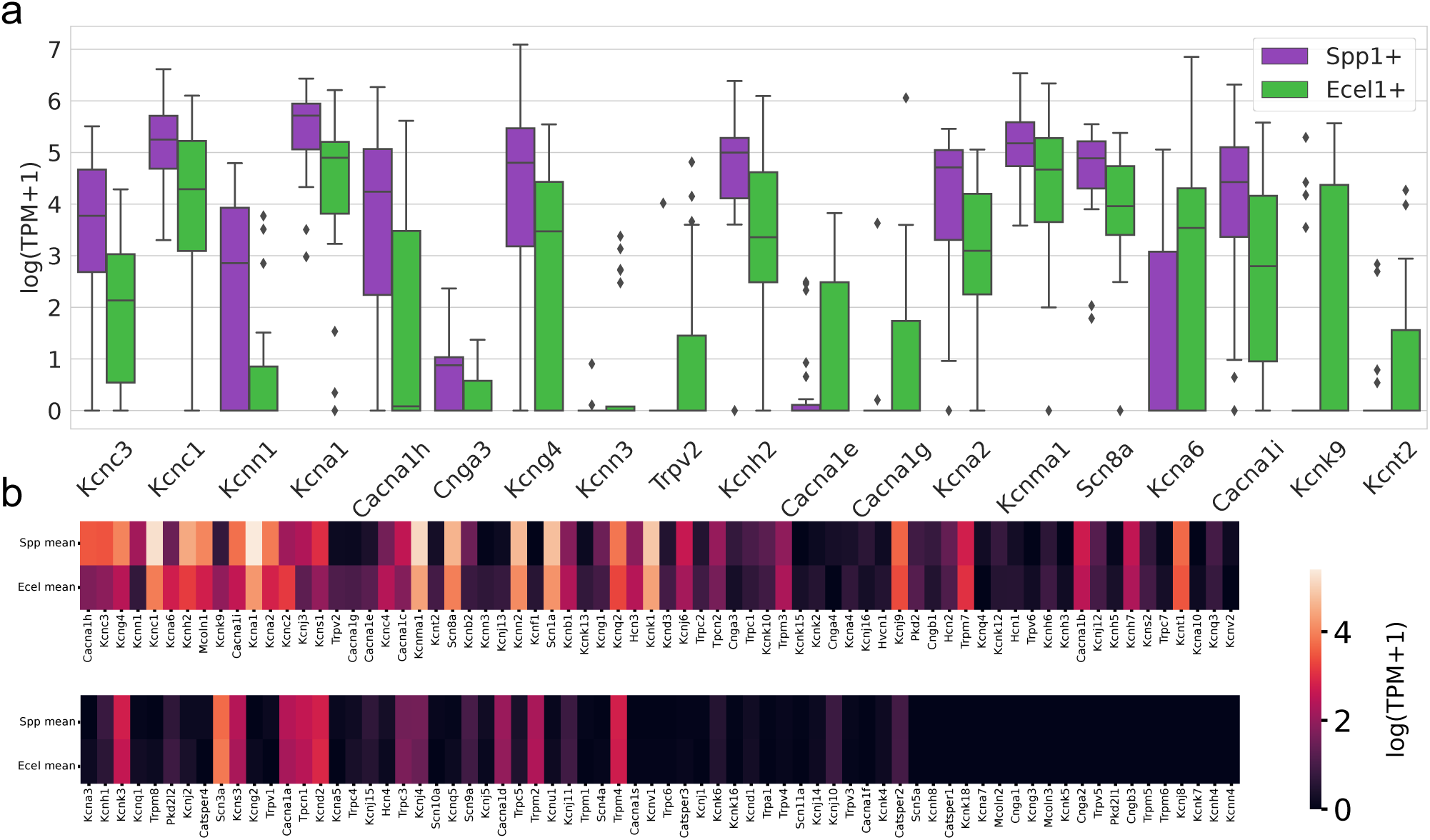
Gene expression level analysis from Li et al. [2020]. **a.** Gene expression levels between Ecel+ and Spp+ cells for genes with the largest differences in expression levels between the two cell types **b.** Average gene expression levels in each subtype, ordered by absolute difference.

